# Effects of habitat destruction on coevolving metacommunities

**DOI:** 10.1101/2021.12.14.472579

**Authors:** Klementyna A. Gawecka, Fernando Pedraza, Jordi Bascompte

## Abstract

Habitat destruction is a growing threat to biodiversity and ecosystem services. The ecological consequences of habitat loss and fragmentation involve reductions in species abundance and even the extinction of species and interactions. However, we do not yet understand how habitat loss can alter the coevolutionary trajectories of the remaining species or how coevolution, in turn, affects their response to habitat loss. To investigate this, we develop a spatially explicit model which couples metacommunity and coevolutionary dynamics. We show that, by changing the size, composition and structure of local networks, habitat destruction increases the diversity of coevolutionary outcomes across the landscape. Furthermore, we show that while coevolution dampens the negative effects of habitat destruction in mutualistic networks, its effects on the persistence of antagonistic communities are less predictable.

## 1 Introduction

Habitat loss and fragmentation are major drivers of the loss of biodiversity (Millennium Ecosystem Assessment, 2005; Pereira et al., 2010; IPBES, 2019). Reduced habitat availability and connectivity decrease species abundance and increase their extinction risk (e.g., Hanski et al., 2013; Chase et al., 2020). As species are embedded in complex networks of interactions, the extinction of one species may trigger further extinctions through the loss of interactions (e.g., Dunne et al., 2002; Memmott et al., 2004; Bascompte et al., 2019), thus exacerbating the negative effects of habitat destruction. Changes in community composition may also reduce their robustness to future habitat loss and fragmentation (Grass et al., 2018). Ultimately, the loss of species and interactions results in the loss of ecosystem function (Dobson et al., 2006; Thompson et al., 2017; Ross et al., 2021) and the services on which we desperately rely on (IPBES, 2019).

While the consequences of habitat destruction on ecological processes have been thoroughly studied (e.g., Evans et al., 2013; Fortuna et al., 2013; McWilliams et al., 2019; Ryser et al., 2019; Heer et al., 2021), its effects on evolutionary processes are not yet fully understood (Hagen et al., 2012; Legrand et al., 2017; Faillace et al., 2021). As ecological changes shape evolutionary outcomes by altering species’ traits, these changes in traits, in turn, affect the ecological processes. This reciprocal interaction between ecology and evolution results in eco-evolutionary feedbacks (Urban et al., 2008; Post and Palkovacs, 2009; Koch et al., 2014; Toju et al., 2017). Furthermore, there is an increasing amount of evidence that evolutionary changes which are ecologically relevant may happen relatively rapidly (Thompson, 1998), such that ecological and evolutionary processes can take place at the same timescales (Hairston et al., 2005; Pelletier et al., 2009; Schoener, 2011). Thus, if habitat destruction affects the ecological processes, it must also influence evolutionary trajectories, and vice versa.

Studying the effects of habitat loss and fragmentation on eco-evolutionary feedbacks calls for the consideration of spatial dynamics. Yet, this can be challenging when approached from either an empirical or a theoretical perspective (Legrand et al., 2017; Gross et al., 2020). Nonetheless, models have been employed to study eco-evolutionary feedbacks in the context of different habitat topologies (Bolchoun et al., 2017; Fronhofer and Altermatt, 2017; Rogge et al., 2019), or the effects of climate change (Norberg et al., 2012; Northfield and Ives, 2013; Åkesson et al., 2021). Additionally, considering entire communities requires the incorporation of the reciprocal evolutionary changes which both interacting and non-interacting species impose on each other, i.e., coevolution. While the interplay between spatial dynamics and coevolution (i.e., the so-called ‘geographic mosaic of coevolution’, Thompson, 2005) has been explored (Gibert et al., 2013; Medeiros et al., 2018; Fernandes et al., 2019), the effects of habitat loss and fragmentation have not been explicitly considered. Therefore, the extent to which habitat destruction influences the eco-(co)evolutionary feedbacks within communities remains largely unknown.

In this study, we develop a spatially explicit model which couples metacommunity and co-evolutionary dynamics. Our model accounts for the effects of local community structure and composition on species’ traits (i.e., the coevolutionary outcome), and how these in turn affect species’ extinction and colonisation rates (Figure 1). First, we investigate the consequences of habitat loss and fragmentation on coevolutionary trajectories by considering changes in trait distributions. We proceed by explaining these effects in terms of changes in network size, composition, and structure. Second, we quantify the role of coevolution in shaping species’ responses to habitat destruction. We also explore the differences between mutualistic and antagonistic networks in coping with habitat loss.

**Figure 1:**
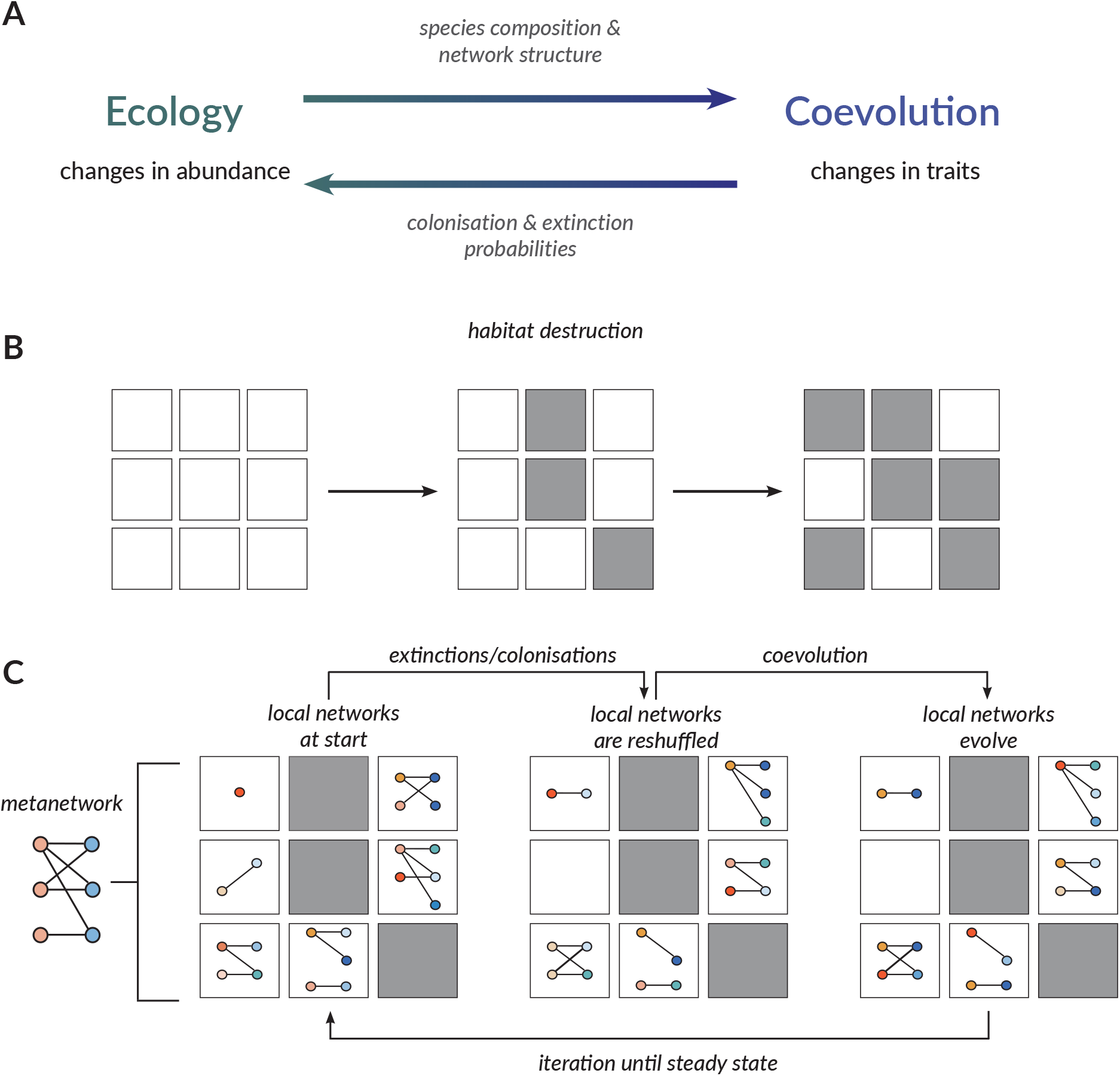
Spatially explicit coevolutionary metacommunity model. (A) Our model includes a feedback loop whereby (1) ecological processes affect coevolutionary dynamics through species composition and local network structure, and (2) the outcome of coevolution (i.e., changes in species’ traits) affects species’ abundance through colonisation and extinction probabilities. (B) Initially, the landscape consists of pristine habitat patches (white squares), but it is progressively destroyed (grey squares) throughout the simulation. (C) We used empirical networks as the metanetworks in the simulations. Each non-destroyed patch can contain a local network which is a subset of the metanetwork. At each habitat destruction step, we simulate metacommunity dynamics (extinctions and colonisations) followed by coevolution within each local network, repeating this sequence until we reach a steady state at the global scale. Extinction and/or colonisation events lead to the reshuffling of the local networks. The subsequent coevolution results in changes in trait values associated with each species in each patch (here, indicated by a change of colour of a node).

## 2 Materials and methods

### 2.1 Coevolutionary metacommunity model

We combined a cellular-automaton-based model for spatial metacommunity dynamics (see Section 2.1.1) with a coevolutionary model for ecological networks (see Section 2.1.2) to simulate the coupled eco-evolutionary (*eco* ↔ *evo*) dynamics in a landscape undergoing habitat destruction (Figure 1). The landscape has dimensions of 100 × 100 identical square patches. We performed simulations with mutualistic (plant-pollinator and seed-disperser) and antagonistic (host-parasite and plant-herbivore, see Table S1) networks from the Web of Life repository (www.web-of-life.es; Fortuna et al., 2014). We used each empirical network as the metanetwork (i.e., the network of all species and interactions across the entire pristine landscape, Figure 1C), and assumed that all patches of pristine habitat are able to harbour the metanetwork.

#### 2.1.1 Spatial dynamics

We adopted a patch dynamics perspective (Leibold et al., 2004) whereby the patches can be occupied or empty, and the spatial matacommunity dynamics are governed by local extinction and colonisation events. While the rules for extinctions and colonisations are based on those proposed by Fortuna et al. (2013) for mutualistic networks, we modified them for the antagonistic networks. Moreover, our rules include the dependence on the degree of trait matching arising from coevolution (i.e., the *eco* ← *evo* coupling, Figure 1A). In other words, species’ extinction and colonisation probabilities depend on the similarity between their traits and the traits of their partners (Hagen et al., 2012). We defined trait matching at time *t* between species *i* and *j* as:

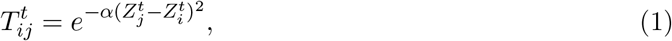

where 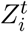 is the mean trait value of the population of species *i* at time *t*, and *α* is a constant controlling the sensitivity of 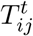 to trait matching (here, varied between 0.02 and 2).

If a species is present in a patch at a given time step, it has a probability of becoming extinct at the next time step. The effective extinction probabilities, 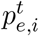, are summarised in Table 1. For both the resource and the consumer in mutualism, as well as the consumer in antagonism, they are independent of the species composition in the patch. Conversely, the effective extinction probability of the resource in antagonism depends on the number of consumer partners present in the patch and their trait matching. In other words, the more consumer partners a resource has, and the higher the trait similarity between them, the more likely it is to become extinct. However, following Andreazzi et al. (2017, 2020), we assume a critical mismatch between trait values of the resource and its antagonistic consumer, *ε*, above which the consumer has no effect on the resource’s extinction probability. Thus, provided that the trait difference is less than or equal to the critical mismatch, the effective extinction probability of the resource increases with the number of consumer partners and the trait similarity between them according to a saturating function. In this study, we varied *ε* between 1 and 10 for all species.

**Table 1:**
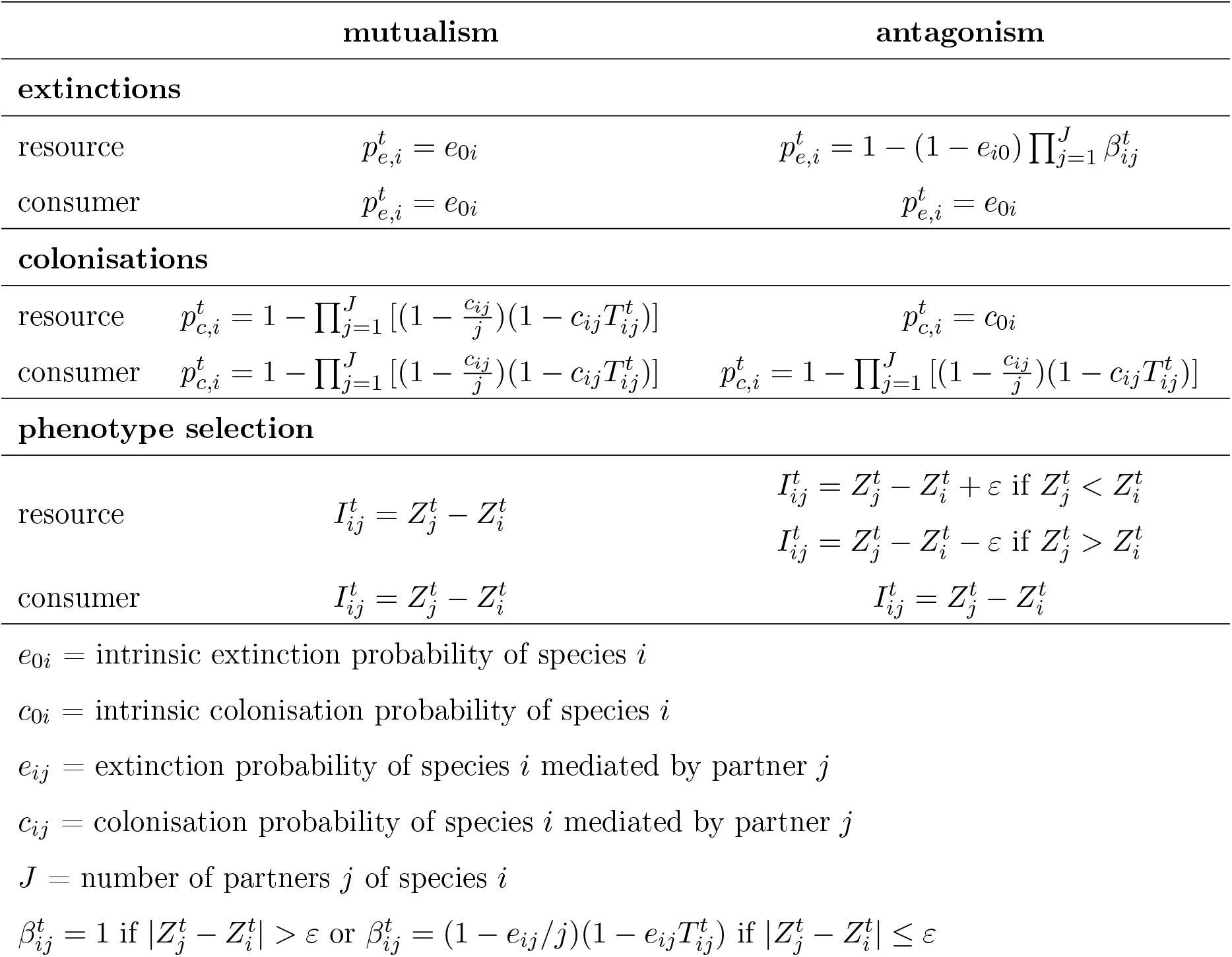
Effective extinction, 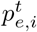, and colonisation, 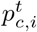, probabilities, and phenotype selection in mutualism and antagonism.

|  | mutualism | antagonism |
| --- | --- | --- |
| **extinctions** |  |  |
| resource | $p_{e,i}^t = e_{0i}$ | $p_{e,i}^t = 1 - (1 - e_{i0}) \prod_{j=1}^J \beta_{ij}^t$ |
| consumer | $p_{e,i}^t = e_{0i}$ | $p_{e,i}^t = e_{0i}$ |
| **colonisations** |  |  |
| resource | $p_{c,i}^t = 1 - \prod_{j=1}^J [(1 - \frac{c_{ij}}{j})(1 - c_{ij}T_{ij}^t)]$ | $p_{c,i}^t = c_{0i}$ |
| consumer | $p_{c,i}^t = 1 - \prod_{j=1}^J [(1 - \frac{c_{ij}}{j})(1 - c_{ij}T_{ij}^t)]$ | $p_{c,i}^t = 1 - \prod_{j=1}^J [(1 - \frac{c_{ij}}{j})(1 - c_{ij}T_{ij}^t)]$ |
| **phenotype selection** |  |  |
| resource | $I_{ij}^t = Z_j^t - Z_i^t$ | $I_{ij}^t = Z_j^t - Z_i^t + \varepsilon$ if $Z_j^t < Z_i^t$<br>$I_{ij}^t = Z_j^t - Z_i^t - \varepsilon$ if $Z_j^t > Z_i^t$ |
| consumer | $I_{ij}^t = Z_j^t - Z_i^t$ | $I_{ij}^t = Z_j^t - Z_i^t$ |

$e_{0i}$ = intrinsic extinction probability of species $i$
$c_{0i}$ = intrinsic colonisation probability of species $i$
$e_{ij}$ = extinction probability of species $i$ mediated by partner $j$
$c_{ij}$ = colonisation probability of species $i$ mediated by partner $j$
$J$ = number of partners $j$ of species $i$
$\beta_{ij}^t = 1$ if $|Z_j^t - Z_i^t| > \varepsilon$ or $\beta_{ij}^t = (1 - e_{ij}/j)(1 - e_{ij}T_{ij}^t)$ if $|Z_j^t - Z_i^t| \leq \varepsilon$

Similarly, a species which is absent from a patch at a given time step has a probability of colonising that patch from one of its eight nearest neighbours (i.e., Moore’s neighbourhood, and assuming a reflective boundary condition) at the next time step. The colonisation events from different patches are considered independent of each other. The effective colonisation probabilities, 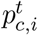, are summarised in Table 1. The consumer (in both mutualism and antagonism) can colonise a patch where at least one of its resource partners is present, thus accounting for the dependence of the consumer on its resources (Fortuna et al., 2013). Conversely, the resource in mutualism can colonise any non-destroyed patch from a neighbouring patch in which both the resource and at least one of its consumer partners are present. This simulates consumer-mediated dispersal (Fortuna et al., 2013). The effective colonisation probability of these species (i.e., the consumers and the mutualistic resource) increases with the number of partners and their trait matching according to a saturating function. Lastly, the resource in antagonism can colonise any non-destroyed patch with a probability that is independent of the species composition in that patch. This simulates a resource that is unable to differentiate between patches with or without its consumers.

The extinction and colonisation probability parameters (i.e., *e*_0*i*_, *c*_0*i*_, *e_ij_*, *c_ij_*) were drawn from uniform distributions for each species (Table S2). Note that the adopted ranges of these values ensure global coexistence of all species in a pristine landscape, although not necessarily in each patch.

#### 2.1.2 Coevolution

Following extinction and colonisation events at a given time step, we simulated the coevolution of the species present in each patch (see Figure 1C), thus incorporating the effect of spatial metacommunity dynamics on coevolution (i.e., the *eco* → *evo* coupling, Figure 1A). We employed the models proposed by Guimarães et al. (2017) and Andreazzi et al. (2017, 2020) for mutualistic and antagonistic networks, respectively. These models incorporate a selection gradient which connects the evolution of trait values with the mean fitness consequences of interactions between species and with the environment. The mean trait evolution of species *i* over a time step is described by:

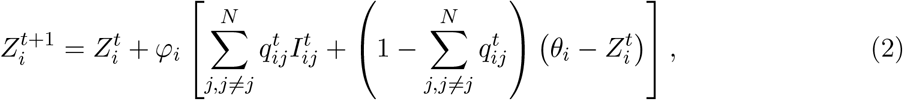

where *φ_i_* controls the slope of the selection gradient and is proportional to the additive genetic variance, *N* is the number of species in the local network, *θ_i_* is the environmental optimum (i.e., the phenotype favoured by the environmental selection), and 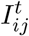 is the phenotype selected by the interaction of species *i* with species *j* (see below). 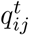 describes the evolutionary effect of species *j* on species *i*, and is defined as:

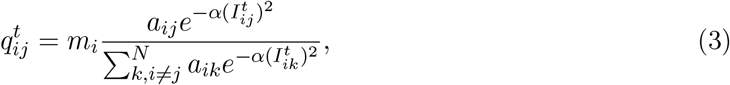

where *m_i_* is the level of coevolutionary selection and a measure of the relative importance of interactions in trait evolution, and *a_ij_* is an element of the binary adjacency matrix, *A*, of the local network (where *a_ij_* = 1 if species *i* and *j* interact, and *a_ij_* = 0 otherwise). The model parameters *φ_i_*, *m_i_* and *θ_i_* for each species are drawn from distributions listed in Table S2, and are assumed to be the same across all patches in the landscape.

Following the approach of Guimarães et al. (2017) and Andreazzi et al. (2017, 2020), we assumed that the phenotype selected by the interaction between two species, 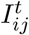, depends on the interaction type. For both the resource and the consumer in mutualism, as well as the consumer in antagonism, selection favours trait matching. Conversely, for the resource in antagonism, selection favours trait mismatch if the trait difference is less or equal to the critical mismatch, *ε* (i.e., if 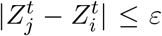). Otherwise (i.e., if 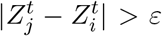), the consumer has no effect on the resource’s fitness. Note that selection can either increase (if 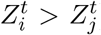) or decrease (if 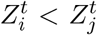) the resource’s trait value. The expressions defining 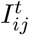 in different scenarios are listed in Table 1.

Additionally, species which do not have partners in a given patch (e.g., due to their partners becoming extinct in the same time step), evolve towards their environmental optima according to:

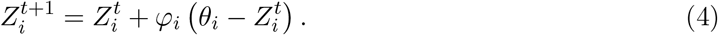

### 2.2 Simulations

Initially, all habitat was pristine and every patch contained the entire metanetwork (i.e., one of the empirical networks in Table S1). We set the initial trait values of all species to their corresponding environmental optima, *θ_i_*. At each time step, we simulated the metacommunity dynamics involving local colonisations and extinctions, followed by coevolution of the species present in each patch (Figure 1C). Thus, we assumed that the ecological and coevolutionary processes take place at the same timescales (Thompson, 1998; Hairston et al., 2005; Pelletier et al., 2009; Schoener, 2011; Hagen et al., 2012). The patches were updated sequentially (in terms of both metacommunity and coevolutionary dynamics), and the model was iterated until a steady state where both the global abundance and the global distribution of trait values of each species no longer changed between time steps. Note that this condition represents a steady state on the landscape scale, but not necessarily within each patch. All results presented here were evaluated at this steady state.

To simulate the destruction of habitat, we then randomly selected 1000 patches (i.e., 10% of the landscape), and changed their state to ‘destroyed’ (Figure 1B). Any species present in these patches immediately became locally extinct, and these patches could not be recolonised. This was followed by iteration over time steps (as described above) until reaching a new steady state. We continued this process until the entire landscape was destroyed.

In order to assess the role of coevolution in species’ response to habitat destruction, we considered two additional scenarios where the species’ trait values were not allowed to change throughout the habitat destruction simulation: ‘evolution’ and ‘frozen coevolution’ (see descriptions in Table 2). First, by comparing these two scenarios, we quantified the differences between evolution and coevolution. Second, by comparing the ‘frozen coevolution’ with ‘continuous co-evolution’ simulations (i.e. where species’ traits are allowed to change as described in Sections 2.1.1 and 2.1.2), we assessed the effect of species being able to readapt to the changing community composition.

**Table 2:**
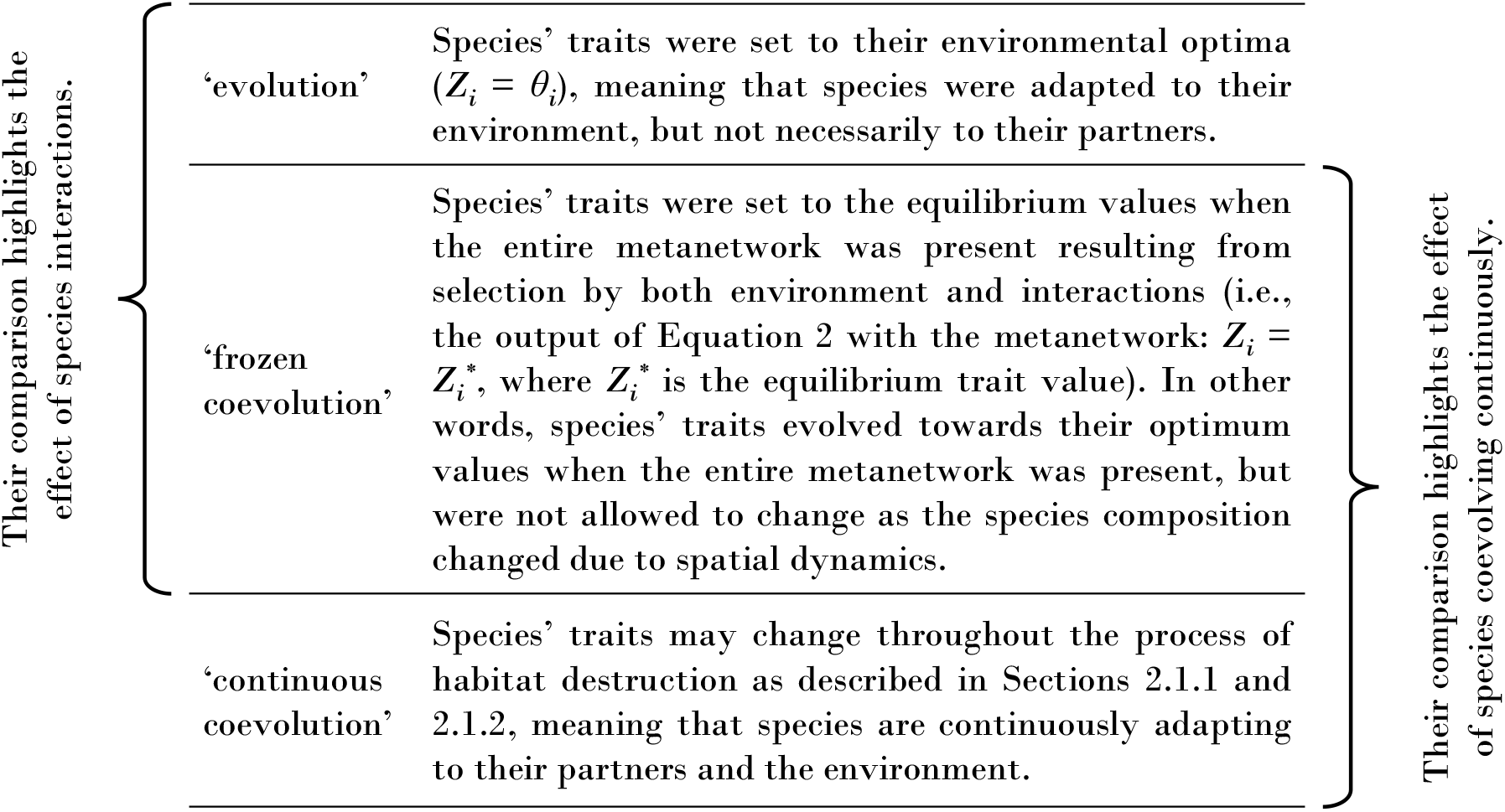
Description of simulation scenarios.

While the simulations described involve destroying habitat patches randomly, we also considered an alternative scenario where habitat was destroyed non-randomly. Starting with 100 randomly selected patches, we subsequently destroyed their adjacent patches, resulting in clusters of destroyed habitat which increased in size. As the results of these simulations show generally similar trends to those with random habitat destruction, they are presented in Supporting Information for brevity (Figures S17-S19).

We performed simulations with 20 networks (10 mutualistic and 10 antagonistic, Table S1). For brevity, here we present the results of one mutualistic (network *M_SD_012*) and one antagonistic network (network *A_PH_004*), whereas the complete set of results can be found in the Supporting Information (Figures S1-S7). Additionally, we investigated the robustness of our results to: the patch removal sequence, the environmental optima (*θ*), the strength of coevolutionary selection (*m*), the trait matching sensitivity (*α*), and the sensitivity of resource to its consumer (*ε*). We found that the former two parameters have a negligible effect, and thus we present only one set of results here. In the case of mutualism, although varying *m* and *α* led to some quantitative differences, the general trends remained the same. Conversely, in antagonistic systems, *m*, *α* and *ε* affected the role of coevolution in shaping species response to habitat loss. Their effects are shown in Figures S8-S10, Figures S11-S13 and S14-S16, respectively, while the results presented here are limited to the intermediate scenarios with mean *m* = 0.7, *α* = 0.2 and *ε* = 5. All simulations were performed with Julia version 1.4.2 (Bezanson et al., 2017), whereas data postprocessing and visualisation was done with R version 3.6.2 (R Core Team, 2020).

### 2.3 Network structure, *β*-diversity and trait matching

We analysed how habitat loss affects ecology by measuring changes in local network structure and composition. Similarly, we quantify the effects of habitat loss on coevolution by measuring changes in species’ trait matching.

For the measure of network structure, we considered network size (the number of species in the local network as a fraction of the number of species in the empirical metanetwork), connectance (the proportion of all possible interactions that are realised in the local network), nestedness and modularity. To quantify nestedness, we adopted the measure proposed by Fortuna et al. (2019), which is equivalent to the NODF metric (Almeida-Neto et al., 2008), but does not penalize the contribution to nestedness of species with the same number of partners. To quantify modularity, we used the ‘igraph’ package in R with a multi-level modularity optimization algorithm (Blondel et al., 2008). For all local networks, we summarised the four descriptors using a principal component analysis (PCA). The first principal component (PC1) explained 69% of the variance and was strongly correlated with connectance (88%), nestedness (93%) and modularity (−94%), whereas the second principal component (PC2) explained 22% of the variance and was strongly correlated with network size (88%).

To assess the differences in species composition across patches, we adopted a *β*-diversity measure. For each fraction of habitat loss, we sampled 100 patches, and calculated the species composition dissimilarity on all their pairwise combinations using the Whittaker measure (Whittaker, 1960). This was done with the ‘betalink’ package in R (Poisot et al., 2012).

Trait matching between interacting species was calculated using Equation 1. We defined network trait matching as the mean trait matching in every local network.

## 3 Results

### 3.1 Effects of habitat destruction on coevolution

We begin by analysing the effects of habitat destruction on the outcome of coevolution (Figure 2). We found that, for the majority of species, trait variability increases with habitat loss, signifying a greater heterogeneity of trait values across the landscape. Comparing the two interaction types, we see that mutualists have a smaller increase in trait variability with habitat loss than antagonists. Moreover, both guilds in mutualism exhibit similar responses. Conversely, while most consumer species in antagonism continuously increase their trait heterogeneity, resource species first increase but then decrease trait diversity as habitat is lost. This is due to the consumers becoming extinct and resources evolving towards their environmental optima, and thus lowering the diversity of trait values across the landscape. We observed similar trends for all simulated mutualistic and antagonistic networks (Figure S1).

**Figure 2:**
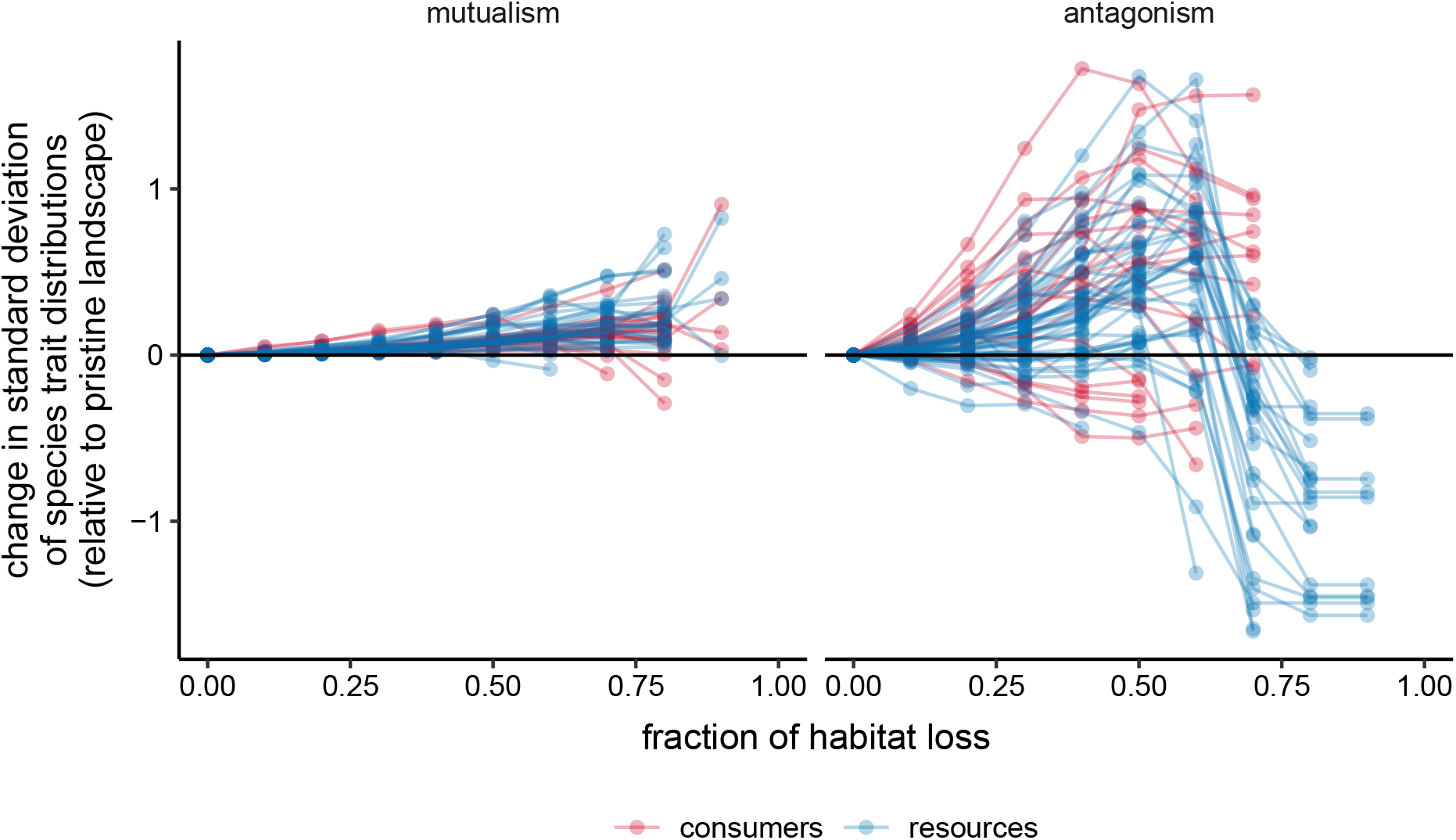
Change in trait variability in the landscape across an increasing fraction of habitat loss in simulations with a mutualistic (left) and an antagonistic (right) network. The change is calculated relative to the standard deviation of trait values in a pristine landscape, such that positive values indicate an increase in trait variability. Each line represents a species.

The above changes in the coevolutionary trajectories can be explained by changes in size, composition and structure of local networks. Firstly, habitat loss reduces species abundance, making the local networks smaller (Figures 3 and S2, bottom panels). Smaller networks achieve higher trait similarity (see Figure 5). Secondly, habitat loss makes the local species composition more varied across the landscape (Figure S3). Since dissimilar networks imply different coevolutioary trajectories, we observe an increase in trait diversity with habitat loss (Figure 2). The changes in network size and composition, and thus also trait variability, are more pronounced in the case of antagonistic communities. Thirdly, the structure of local networks is altered as habitat is destroyed (Figures 3 and S2, top panels). While mutualistic networks become more connected and nested (positive PC1 values), antagonistic networks become more modular (negative PC1 values). As a consequence, we find larger trait variability in antagonistic than mutualistic communities, since nested networks achieve greater trait similarity across all species (Pedraza et al., 2021), whereas higher modularity enables species to evolve greater trait matching within their modules but not necessarily across modules.

**Figure 3:**
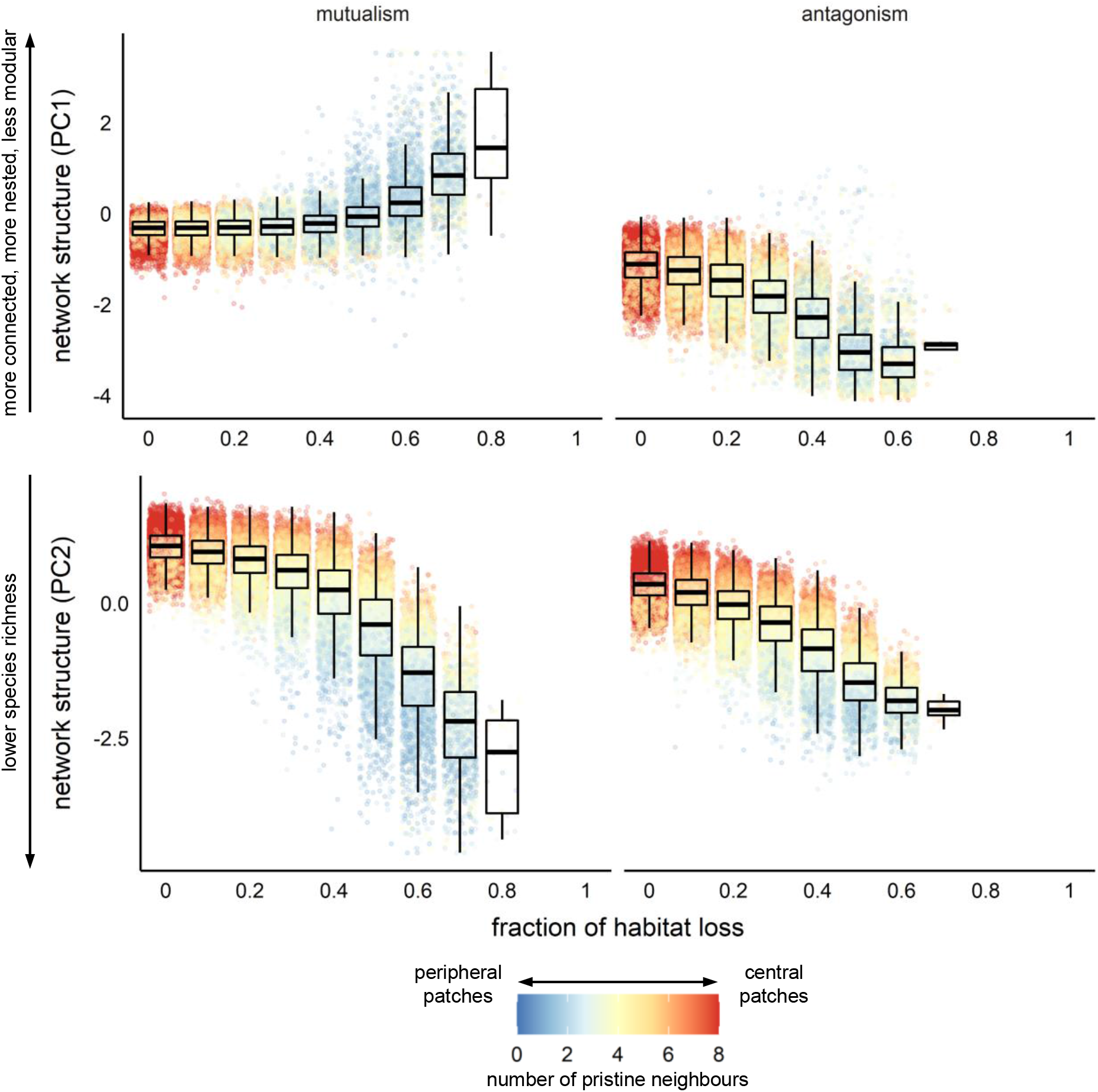
Effect of habitat loss on our measures of network structure of one mutualistic (left) and one antagonistic (right) network. Each point corresponds to the local network inside each patch. The colours indicate the number of adjacent pristine patches (i.e., between 0 and 8). PC1 explained 69% of variance and was strongly correlated with connectance, nestedness and modularity, whereas PC2 explained 22% of variance and was strongly correlated with network size.

The above changes in network structure with habitat loss can be explained by considering the order of species extinctions (Figure S4). In mutualism, the extinction thresholds of both consumer and resource species increase with their degree, meaning that specialists are less abundant and become extinct first, resulting in more nested structures (Figure S5). In antagonism, while the extinction threshold of consumers is pushed towards higher values of habitat loss with their degree, the opposite is true for resources. This sequence of species extinctions results in increased modularity of local networks (Figure S5).

Finally, it is worth noting that peripheral patches (i.e., those with fewer adjacent pristine patches) tend to harbour smaller, more connected and nested mutualistic networks, or smaller and more modular antagonistic networks, than central patches. The exception to this pattern occurs in highly fragmented landscapes with antagonistic interactions, where peripheral patches contain smaller but less modular networks than central patches. In this case, the peripheral networks are likely to be so small that they are partitioned in only few modules.

### 3.2 Effects of coevolution on the community’s response to habitat destruction

Once we have looked at how habitat loss affects the coevolutionary dynamics of the resulting species, we next investigate how coevolution affects the response to these metacommunities to habitat loss. To do this, we compared the results of simulations where species’ traits were fixed (the ‘evolution’ and ‘frozen coevolution’ scenarios, see Table 2) with those obtained when they were allowed to change (‘continuous coevolution’ scenario, i.e., results discussed so far) by considering the differences in the regional abundance of interactions, i.e., the fraction of patches where a given interaction is present (Figures 4 and S6).

**Figure 4:**
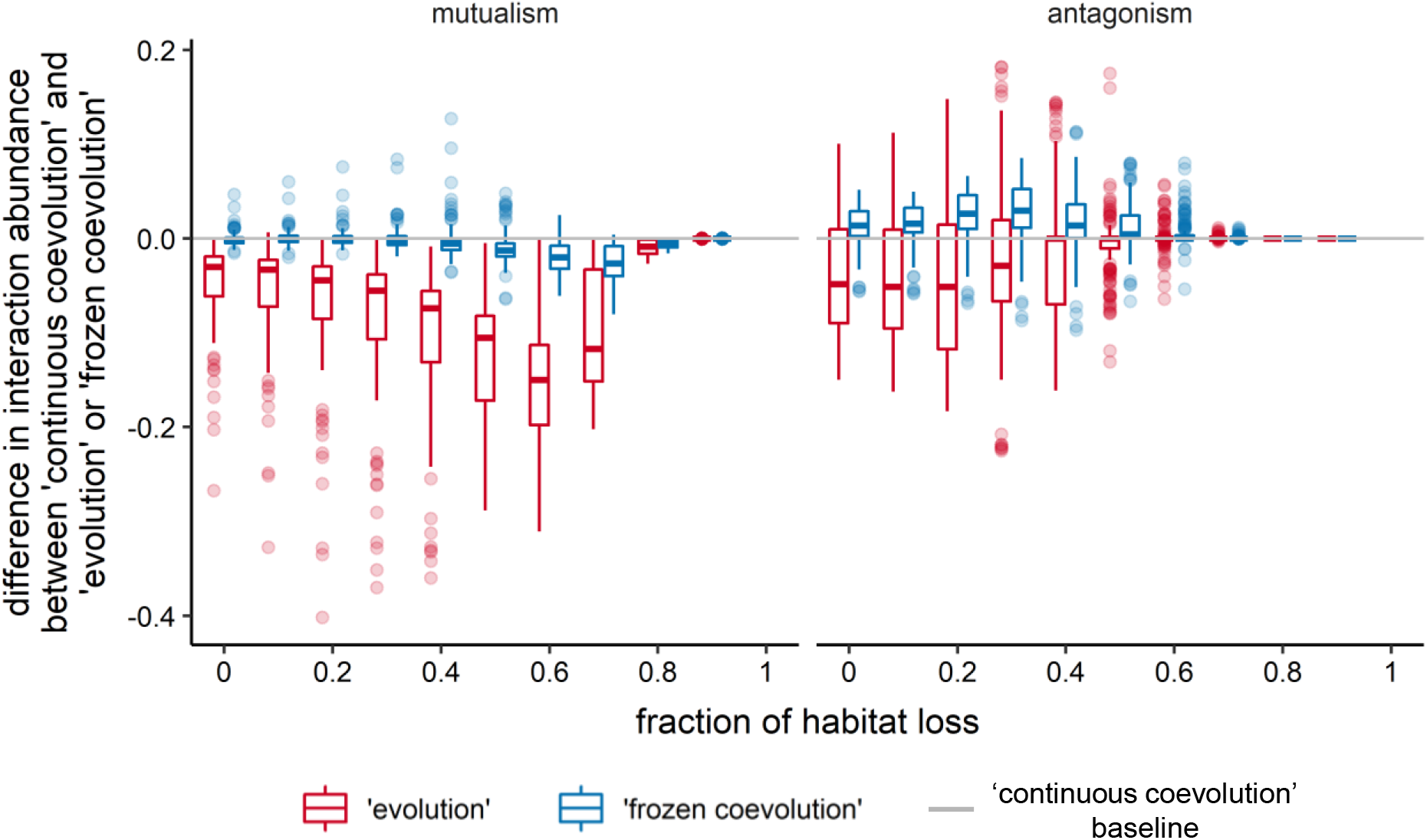
Relative abundance of each interaction in a mutualistic (left) and an antagonistic (right) network as habitat is progressively destroyed. The figure compares the interaction abundance in ‘evolution’ and ‘frozen coevolution’ scenarios in relation to the ‘continuous coevolution’ baseline. Abundance was calculated as the number of patches harbouring each interaction as a fraction of non-destroyed patches in the landscape. Positive values indicate that the abundance in ‘evolution’ or ‘frozen coevolution’ simulations is greater than in the ‘continuous coevolution’ simulations, at the same fraction of habitat loss. In the ‘evolution’ simulations, species’ traits are set to their environmental optima. In the ‘frozen coevolution’ simulations, species’ traits are set to their optimum values when the entire metanetwork is present (see Table 2).

Firstly, comparing the ‘evolution’ and ‘frozen coevolution’ simulations shows the effect of species interactions in shaping traits. We found that, in the case of mutualism, the ‘frozen coevolution’ simulations yielded higher interaction regional abundances than the ‘evolution’ simulations. This means that, allowing trait changes to be driven by interactions, rather than purely by the environment, dampens the negative effects of habitat loss. However, in antagonism, the differences between ‘evolution’ and ‘frozen coevolution’ are generally smaller. Moreover, some networks achieve higher interaction abundances with ‘frozen coevolution’, while others with ‘evolution’ (see Figure S6).

Secondly, comparing ‘frozen coevolution’ and ‘continuous coevolution’ simulations demonstrates the effect of species being able to coevolve continuously throughout the habitat destruction process. At low fractions of habitat loss, there is little difference between the abundance of mutualistic interactions in the two scenarios, with some interactions benefiting from (i.e., those below the zero line in Figure 4) and some being hindered by (i.e., those above the zero line in Figure 4) continuous coevolution. However, in fragmented environments, abundance of mutualistic interactions is higher in the ‘continuous coevolution’ simulations, meaning that the ability to readapt to new community compositions aids the species response to habitat loss. Conversely, in antagonism, some networks achieve lower and others higher interaction abundances in the ‘continuous coevolution’ simulations. Moreover, the effect of coevolution on antagonistic interaction abundance depends not only on the network, but also on the strength of coevolutionary selection (*m*, Figure S9), the trait matching sensitivity (*α*, Figure S12), and the sensitivity of resource to its consumers (*ε*, Figure S15). This again demonstrates the unpredictability of the coevolutionary outcomes in antagonistic networks, and suggests that continuous coevolution does not always benefit the response of antagonistic systems to habitat destruction.

Since we assume that species’ colonisation and extinction probabilities depend on the trait similarity with their partners, the differences in interaction abundance between the simulations outlined above are related to trait matching (Figures 5 and S7). In mutualism, colonisation probabilities of both guilds increase with trait matching. Therefore, higher interaction abundances in the ‘frozen coevolution’ than ‘evolution’ simulations result from coevolution leading to higher trait matching than would be expected if species evolved to their environmental optima (i.e., compare red and blue dashed lines in Figures 5 and S7). Comparing ‘frozen coevolution’ simulations (blue dashed lines in Figures 5 and S7) with ‘continuous coevolution’ (box plots in Figures 5 and S7), we see that, at high fractions of habitat loss, trait matching tends to be higher in the latter. This explains the higher interaction abundance in the ‘continuous coevolution’ simulations. However, these relations are reversed in antagonism due to the conflicting interests of the two guilds. While higher trait matching increases consumer’s colonisation probability, it also increases resource’s extinction chances. Overall, these conflicting forces result in higher trait matching translating to lower interaction abundance (compare abundances in Figures 4 and S6 with trait matching in Figures 5 and S7 for any type of simulation).

**Figure 5:**
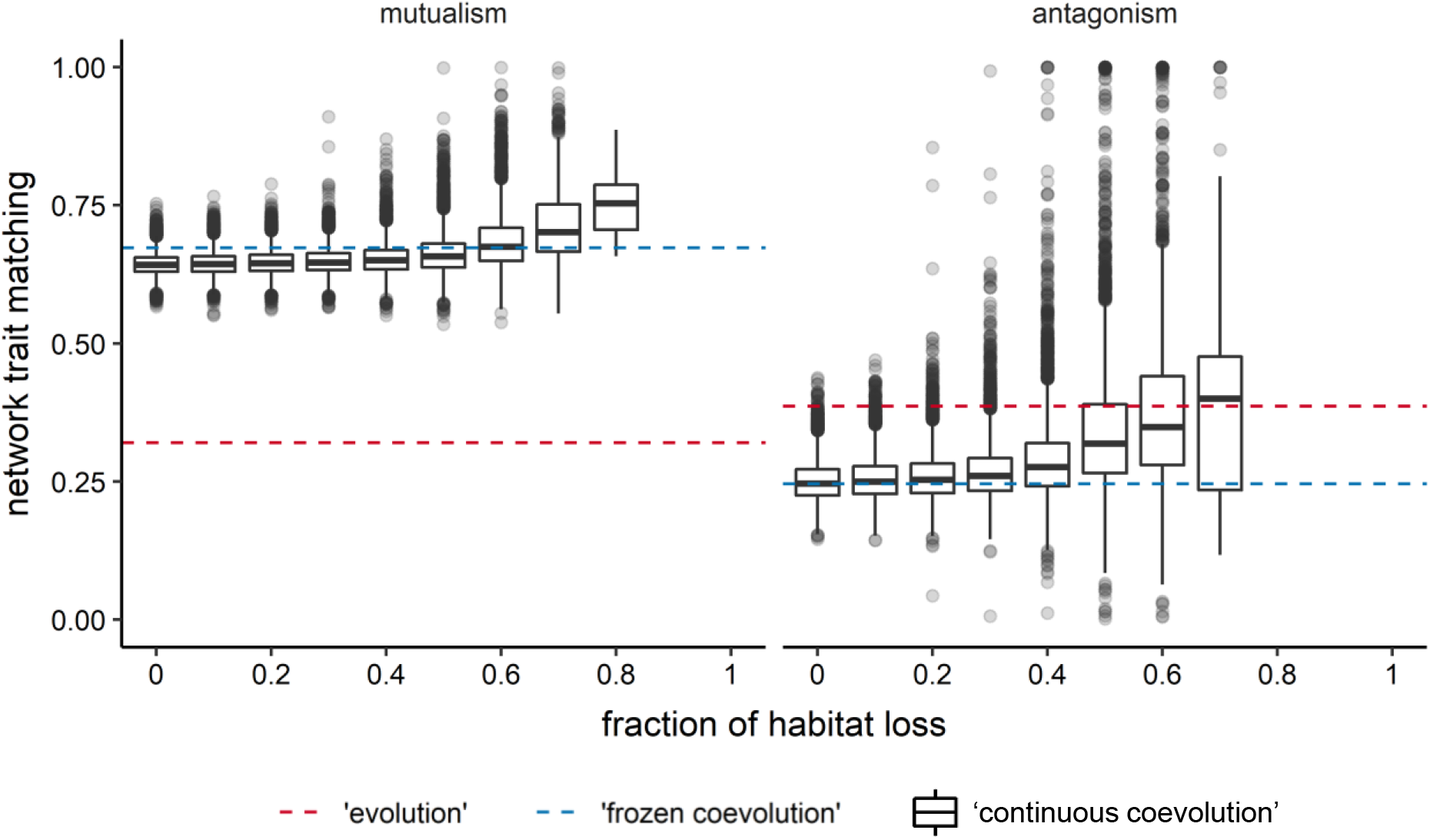
Network trait matching across a gradient of habitat destruction in simulations with a mutualistic (left) and an antagonistic (right) network. Box plots summarise trait matching in all local networks in the landscape at each fraction of habitat loss in ‘continuous coevolution’ simulations. The red dashed horizontal lines correspond to the mean matching of environmental optima of all interacting species (‘evolution’ simulations). The blue dashed horizontal lines indicate the mean matching of optimum trait values when the entire empirical metanetwork is present (‘frozen coevolution’ simulations).

## 4 Discussion

Our spatially explicit model, which couples metacommunity and coevolutionary dynamics, enabled us to study (1) the effects of habitat destruction on the coevolutionary processes, and (2) the effects of coevolution on the species’ response to habitat destruction. First, we show that habitat loss leads to changes in the coevolutionary trajectories by altering the size, composition and structure of local networks (Figure 6, top panels). Second, we demonstrate that, while coevolution buffers the negative consequences of habitat loss on mutualists, it may have the opposite effects in the case of antagonistic interactions (Figure 6, bottom panels). Our results broaden the understanding of the eco-evolutionary community dynamics in fragmented landscapes.

**Figure 6:**
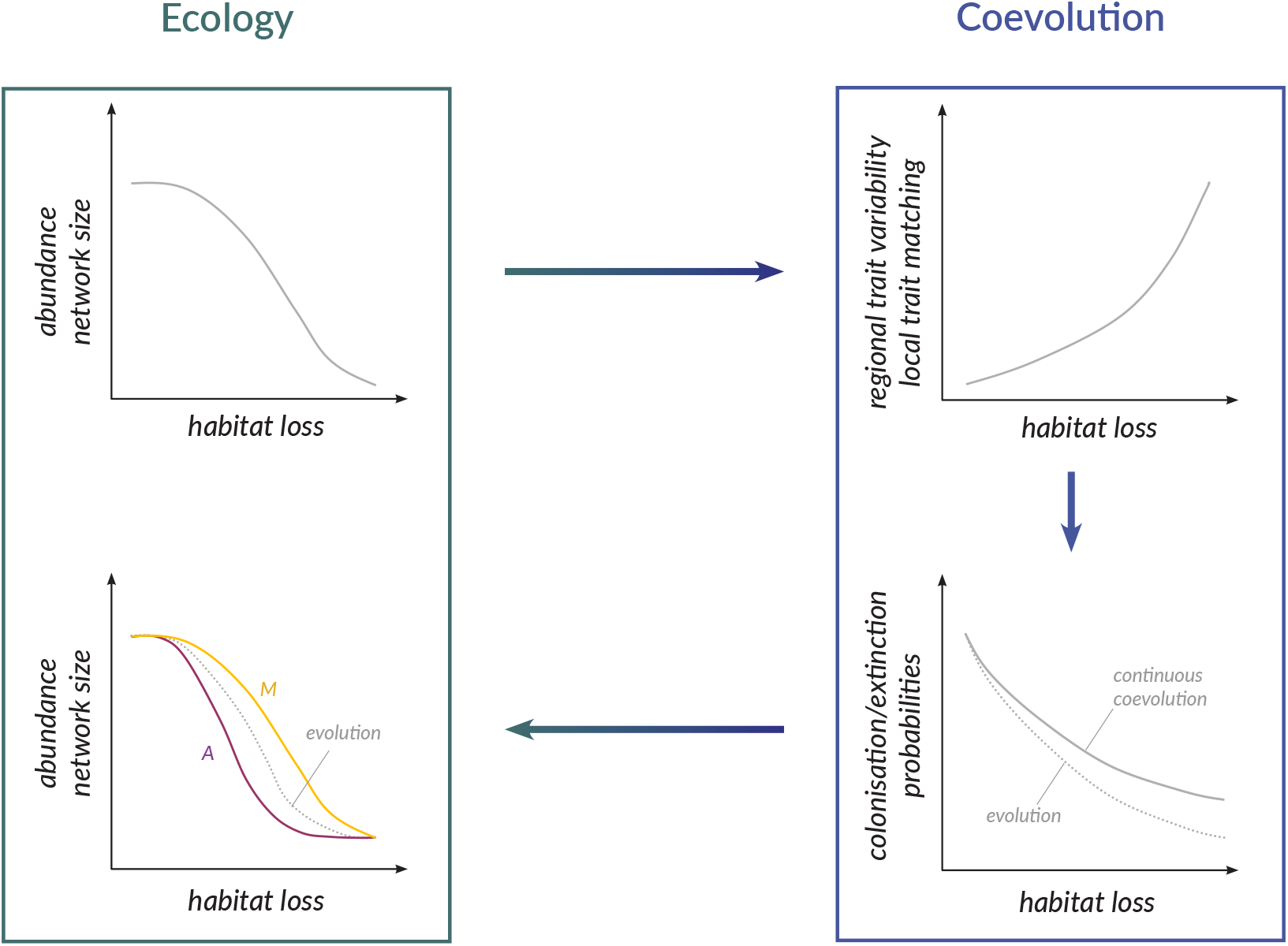
Schematic summary of our results. Habitat destruction reduces species and interaction abundance and local network size (top left panel). This leads to changes in coevolutionary outcomes – an increase in trait variability across the landscape and trait matching locally (top right panel). As habitat is lost, colonisation and extinction probabilities are reduced due to decreasing number of partners (bottom right panel). However, in the case of ‘continuous coevolution’, this reduction is partly offset by the increase in trait matching. The changes in colonisation and extinction probabilities manifest themselves in changes in species and interaction abundance (bottom left panel), which are beneficial to mutualistic communities, but can be disadvantageous to antagonistic communities.

Habitat destruction changes network size and composition, thus affecting the local coevolutionary dynamics. The reduction in abundance, and eventual loss of species due to habitat destruction results in the remaining pristine habitat patches harbouring smaller and more densely connected networks (Figures 3 and S2). This has also been observed in empirical studies of plant-pollinator communities (Spiesman and Inouye, 2013) and food webs (Start et al., 2020). While smaller networks achieve higher trait similarity locally (Figures 5 and S7; see also Pedraza et al., 2021), the differences in the coevolutionary outcomes between patches become more pronounced with habitat loss, leading to larger trait variability across the landscape (Figures 2 and S1). This increased spatial heterogeneity of species’ traits is a result of greater variability of local species compositions (Figure S3). Antagonistic networks exhibit larger changes in trait variability with habitat destruction, suggesting that they are more sensitive to the effects of habitat loss than mutualists. These coevolutionary changes may have implications for restoration and/or species translocations. For example, greater trait variability across the landscape means that more populations may lack a shared coevolutionary history, and thus may be unable to interact once reconnected. Our results suggest that this may be more problematic for antagonistic than mutualistic communities.

Furthermore, species loss alters the local network architecture leading to more nested structures in the case of mutualism, but more modular antagonistic networks. While theory suggests that these structural changes may increase persistence (Thébault and Fontaine, 2010; Stouffer and Bascompte, 2011; Rohr et al., 2014), it is important to remember that this comes at the cost of biodiversity loss in the context of habitat destruction. Our results also show that the periphery of pristine habitat fragments is more susceptible to these changes in network size and structure, and thus, should be prioritised for conservation.

The alterations to coevolutionary processes driven by habitat loss and fragmentation must also influence the species’ ecological response through eco-evolutionary feedbacks (Urban et al., 2008; Post and Palkovacs, 2009; Koch et al., 2014). To quantify this hidden effect, we considered alternative scenarios where species’ traits did not change throughout habitat destruction (Table 2). We asked: (1) whether it is better to coevolve with partners or simply evolve when dealing with habitat loss, and (2) whether the continuous ability to readapt to the changing community composition is beneficial when coping with habitat loss. We found that the answers to these questions depend on the interaction type.

In mutualism, coevolution results in substantially higher interaction abundances than when selection is driven solely by the environment (Figures 4 and S6, ‘evolution’ simulations). Since coevolution selects for trait similarity in both guilds, thus allowing species to adapt better to their partners, their dispersal and/or survival chances increase. Moreover, the ability to coevolve continuously, rather than having traits fixed to values which are optimum in pristine landscapes, provides additional benefits in highly fragmented landscapes (Figures 4 and S6, ‘frozen coevolution’ simulations). In other words, continuous coevolution offsets some of the negative effects of habitat loss on mutualistic communities, thus acting as a negative feedback mechanism.

In the case of antagonistic interactions, the differences between evolution and coevolution are smaller than in mutualism. Also, the highest interaction abundances are achieved with either evolution or coevolution (‘frozen’ or ‘continuous’) depending on the network (Figure S6), the strength of coevolutionary selection (*m*, Figure S9), the trait matching sensitivity (*α*, Figure S12), and the sensitivity of the resource to its consumers (*ε*, Figure S15). In other words, coevolution may be beneficial when coping with habitat loss in some circumstances, but it can also exacerbate its effects in others. This high uncertainty in the coevolutionary outcome is due to the asymmetry between the guilds (where consumers favour trait convergence but resources are selected for trait divergence) creating highly complex eco-evolutionary dynamics. These findings again demonstrate the higher vulnerability, and unpredictability of the response to habitat loss of antagonistic communities.

Previous studies on ecological consequences of habitat loss have shown that destroying patches in clusters, rather than at random, reduces the negative effects of fragmentation and substantially shifts species’ extinction thresholds towards higher values of habitat loss (e.g., Dytham, 1994; Hill and Caswell, 1999; With and King, 1999; Ovaskainen et al., 2002; Gawecka and Bascompte, 2021). Our results confirm that this holds for interaction networks (both mutualistic and antagonistic, see Supplementray Data). Furthermore, we found that, since the local network structure changes less with habitat loss than when destruction is random, the coevolutionary trajectories are also affected to a lesser extent (Figures S18-S17). This then leads to smaller effects of coevolution on the species’ response to habitat loss (Figure S19). This may also be true in networks with long distance dispersal, as the ability to colonise habitats further away (rather than the nearest patches only as in this study) has been shown to increase the extinction thresholds (Gawecka and Bascompte, 2021). Finally, we postulate that network rewiring (which has not been considered here for simplicity) may further buffer the effects of habitat loss on both the ecological and coevolutionary dynamics, as the flexibility of interaction partners may increase species’ persistence (e.g., Kaiser-Bunbury et al., 2010; Ramos-Jiliberto et al., 2012; Tylianakis and Morris, 2017; Santos et al., 2021).

In summary, by reducing species’ abundance and changing local network structure, habitat destruction increases the trait similarity between partners locally and the heterogeneity of species’ traits across the landscape (Figure 6, top panels). These effects are stronger in antagonistic than mutualistic communities. Furthermore, our study demonstrates that coevolution shapes species’ traits in a way that aids the response of mutualists to habitat loss, but it may be either advantageous or disadvantageous to antagonistic interactions depending on the network and the coevolutionary dynamics within it (Figure 6, bottom panels). This complexity and unpredictability of antagonistic community dynamics creates potential challenges for their conservation or restoration. With habitat destruction continuing to pose a great threat to global biodiversity, a better understanding of both its ecological and evolutionary consequences is of utmost importance.

## Acknowledgements

We thank the members of Bascompte Lab for discussions. Funding was provided by SNSF (grant number 310030_197201 to JB), and the University of Zurich Research Priority Program Global Change and Biodiversity (URPP GCB).

## Data availability

All code used in this study is available on GitHub (https://github.com/kgawecka/habiat_destruction_coevolution).

## Authors’ contributions

KAG, FP and JB designed the study. KAG and FP developed the models, performed the simulations and analysed the data. KAG wrote the first draft of the manuscript, and all authors contributed substantially to revisions.

## Notes

### Competing Interest Statement

The authors have declared no competing interest.

