## Supporting Information for "Effects of habitat destruction on coevolving metacommunities"

### Network data and model parameters

Table S1: Networks simulated in this study (PL - plant-pollinator, SD - plant-seed disperser, HP - host-parasite, PH - plant-herbivore).

| network | resources | consumers | interactions | connectance | nestedness | modularity | reference |
|---|---|---|---|---|---|---|---|
| mutualistic | | | | | | | |
| M_PL_006 | 17 | 61 | 146 | 0.14 | 0.65 | 0.38 | Dicks et al. (2002) |
| M_PL_010 | 31 | 76 | 456 | 0.19 | 0.36 | 0.23 | Elberling & Olesen (unpublished) |
| M_PL_036 | 10 | 12 | 30 | 0.25 | 0.41 | 0.42 | Olesen (unpublished) |
| M_PL_037 | 10 | 40 | 72 | 0.18 | 0.29 | 0.48 | Montero (2005) |
| M_PL_059 | 13 | 13 | 71 | 0.42 | 0.93 | 0.20 | Bezerra et al. (2009) |
| M_SD_005 | 25 | 13 | 49 | 0.15 | 0.36 | 0.52 | Carlo et al. (2003) |
| M_SD_008 | 16 | 10 | 110 | 0.69 | 0.86 | 0.10 | Frost (1980) |
| M_SD_010 | 50 | 14 | 234 | 0.33 | 0.53 | 0.21 | Snow and Snow (1971) |
| M_SD_012 | 35 | 29 | 146 | 0.14 | 0.37 | 0.36 | Galetti and Pizo (1996) |
| M_SD_025 | 7 | 6 | 22 | 0.52 | 0.78 | 0.19 | Sorensen (1981) |
| antagonistic | | | | | | | |
| A_HP_005 | 7 | 13 | 51 | 0.56 | 0.92 | 0.12 | Hadfield et al. (2014) |
| A_HP_008 | 8 | 24 | 37 | 0.19 | 0.42 | 0.47 | Hadfield et al. (2014) |
| A_HP_015 | 3 | 7 | 12 | 0.57 | 1.00 | 0.21 | Hadfield et al. (2014) |
| A_HP_028 | 4 | 15 | 19 | 0.32 | 0.54 | 0.45 | Hadfield et al. (2014) |
| A_HP_032 | 14 | 13 | 32 | 0.18 | 0.19 | 0.48 | Hadfield et al. (2014) |

| | | | | | | | |
|---|---|---|---|---|---|---|---|
| A_HP_035 | 6 | 7 | 15 | 0.36 | 0.66 | 0.34 | Hadfield et al. (2014) |
| A_HP_042 | 21 | 32 | 84 | 0.13 | 0.24 | 0.45 | Hadfield et al. (2014) |
| A_HP_050 | 27 | 35 | 226 | 0.24 | 0.42 | 0.25 | Hadfield et al. (2014) |
| A_PH_004 | 52 | 22 | 184 | 0.16 | 0.33 | 0.40 | Joern (1979) |
| A_PH_005 | 54 | 24 | 173 | 0.13 | 0.34 | 0.40 | Joern (1979) |

Table S2: Model parameters and their values.

| | **mutualism** | **antagonism** |
|---|---|---|
| $e_{0i}$ | $\mathcal{U}(0.14, 0.16)$ | $\mathcal{U}(0.14, 0.16)$ |
| $c_{0i}$ | $\mathcal{U}(0.04, 0.06)$ | $\mathcal{U}(0.14, 0.16)$ |
| $e_{ij}$ | $\mathcal{U}(0.14, 0.16)$ | $\mathcal{U}(0.14, 0.16)$ |
| $c_{ij}$ | $\mathcal{U}(0.04, 0.06)$ | $\mathcal{U}(0.14, 0.16)$ |
| $\alpha$ | 0.02, 0.2, 2 | 0.02, 0.2, 2 |
| $\varepsilon$ | – | 1, 5, 10 |
| $\varphi_i$ | $\mathcal{N}(0.5, 0.01)$ | $\mathcal{N}(0.5, 0.01)$ |
| | $\mathcal{N}(0.5, 0.01)$ | $\mathcal{N}(0.5, 0.01)$ |
| $m_i$ | $\mathcal{N}(0.7, 0.01)$ | $\mathcal{N}(0.7, 0.01)$ |
| | $\mathcal{N}(0.9, 0.01)$ | $\mathcal{N}(0.9, 0.01)$ |
| $\theta_i$ | $\mathcal{U}(0, 10)$ | $\mathcal{U}(0, 10)$ |

### Random habitat destruction

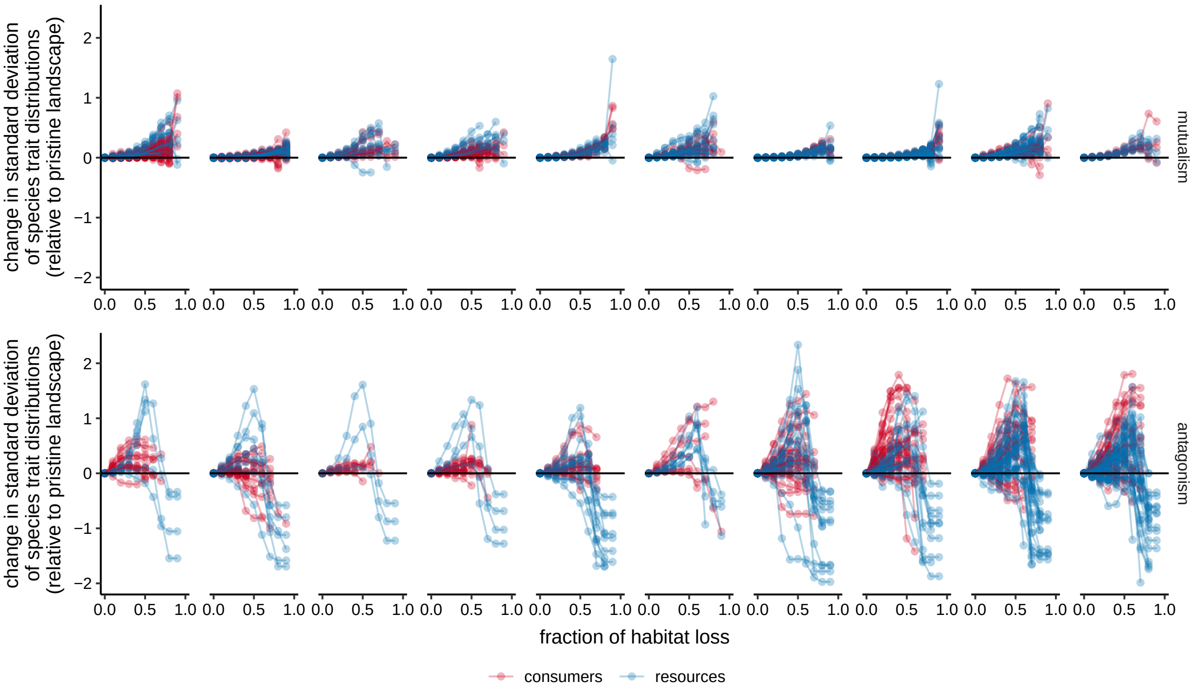


Figure S1: Change in trait variability in the landscape across an increasing fraction of habitat loss in simulations with mutualistic (top) and antagonistic (bottom) networks. The change is calculated relative to the standard deviation of trait values in a pristine landscape, such that positive values indicate an increase in trait variability. Each line represents a species. Panels correspond to different networks in the same order as listed in Table S1.

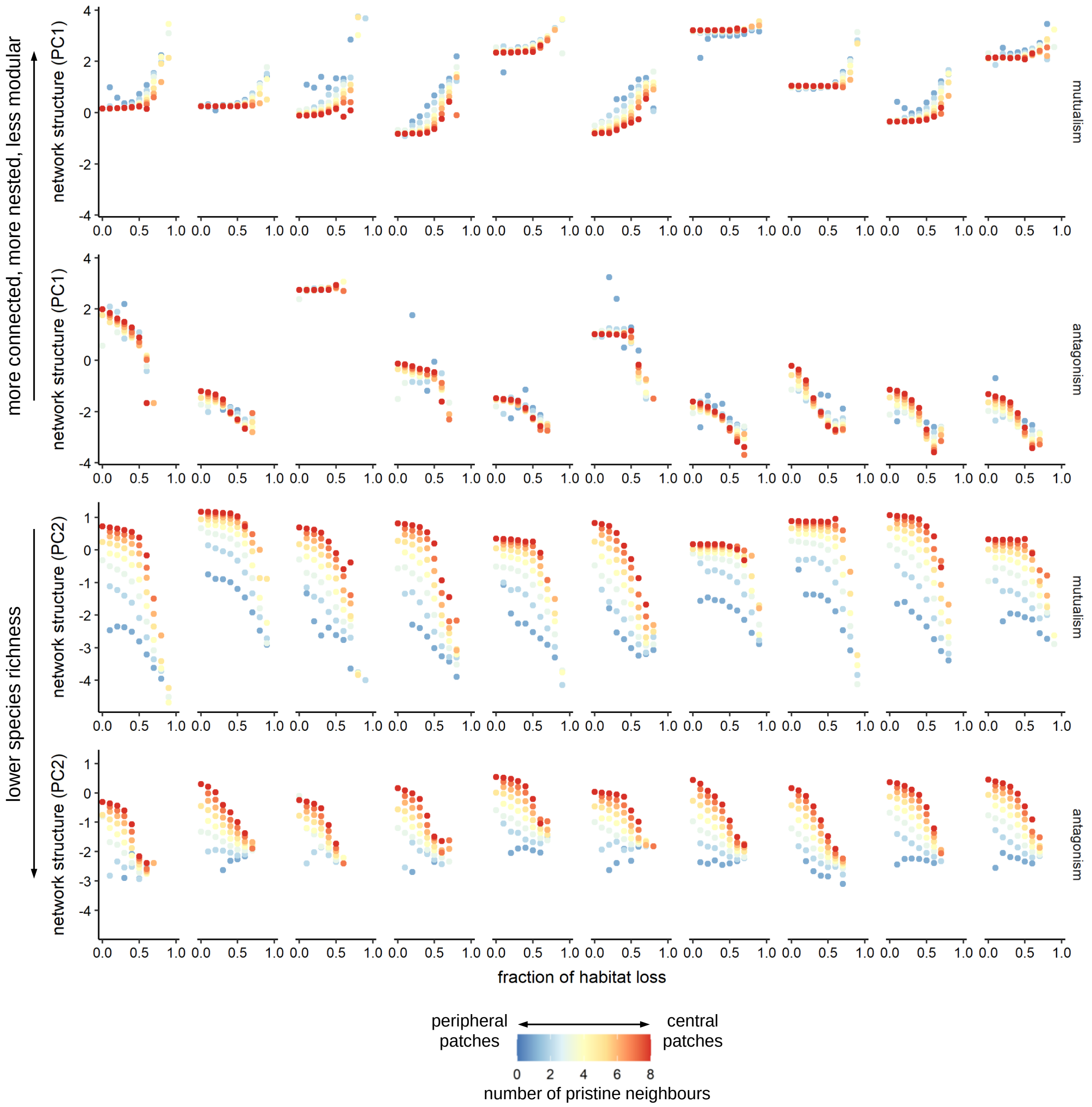


Figure S2: Effect of habitat loss on our measure of network structure of mutualistic and antagonistic networks. Panels correspond to different networks in the same order as listed in Table S1. Each point represents the mean network structure across all patches with the same number of pristine neighbours. The colours indicate the number of neighbouring pristine patches (i.e., between 0 and 8). PC1 explained 69% of variance and was strongly correlated with connectance, nestedness and modularity, whereas PC2 explained 22% of variance and was strongly correlated with network size.

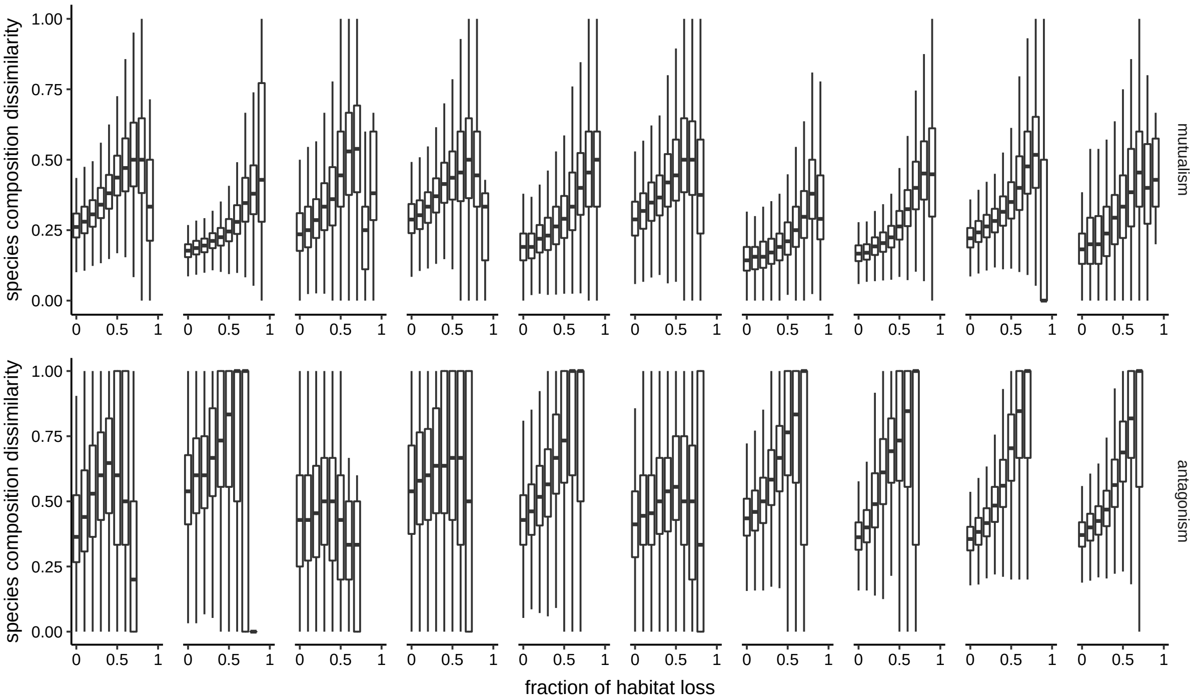


Figure S3: Effect of habitat loss on dissimilarity in local species composition in mutualistic (top) and antagonistic (bottom) networks. Dissimilarity was calculated on all pairwise combinations of 100 sampled patches (per fraction of habitat loss) using the Whittaker $\beta$-diversity measure (Whittaker, 1960). Panels correspond to different networks in the same order as listed in Table S1.

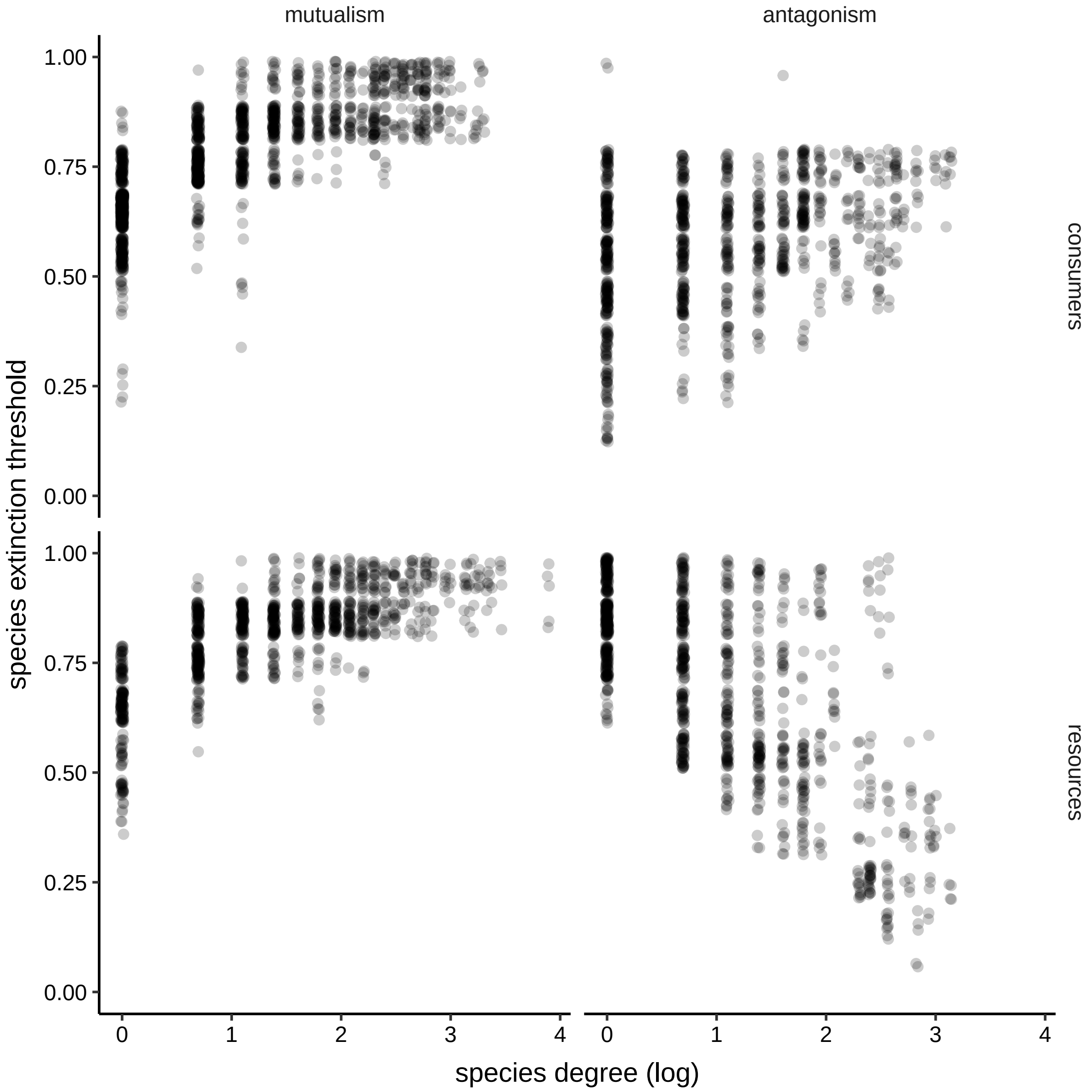


Figure S4: Effect of species degree on their extinction threshold all species in all simulated networks. Species degree is defined as the number of links of each species, and extinction threshold is the fraction of habitat loss at which the species becomes extinct.

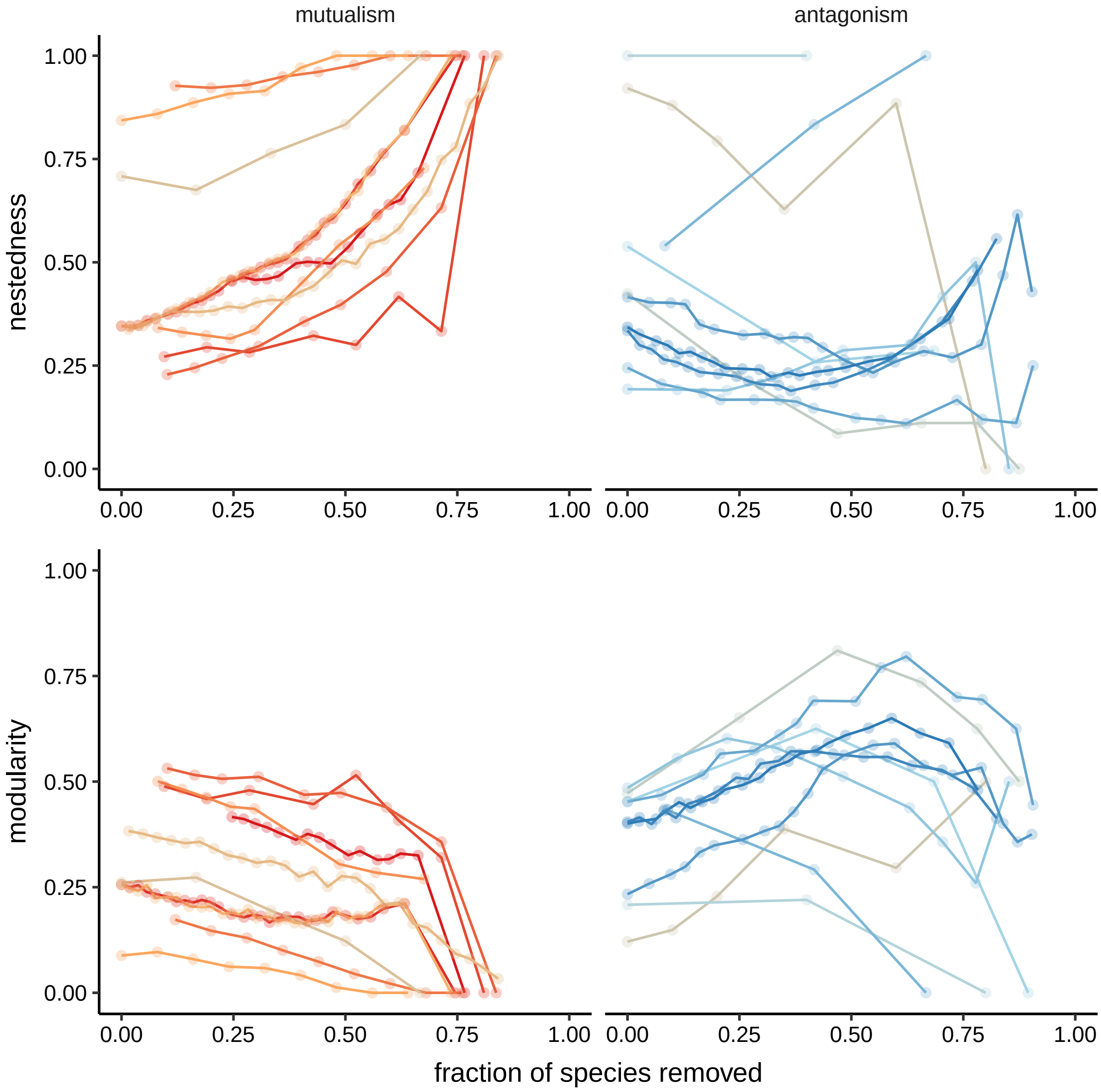


Figure S5: Effect of progressively removing species from mutualistic (left) and antagonistic (right) networks on their nestedness (top) and modularity (bottom). For each empirical network (Table S1), we removed species sequentially following the order of their extinction shown in Figure S4. In case of mutualistic networks, this was done from most to least specialist species (for both guilds). In the case of antagonistic networks, consumers were removed from most to least specialist, whereas resources were removed from most to least generalist. Every time we removed a species, we recalculated nestedness and modularity of the network (points). Each network is indicated by a different colour line.

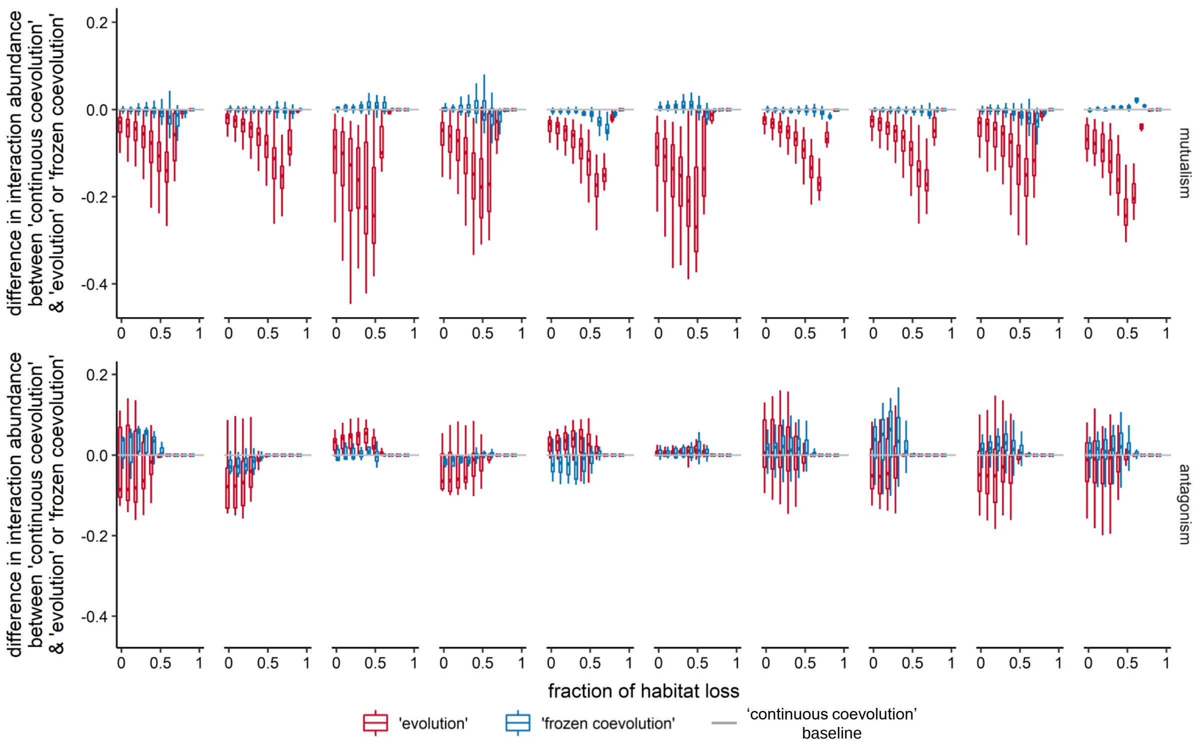


Figure S6: Relative abundance of each interaction in mutualistic (top) and antagonistic (bottom) networks as habitat is progressively destroyed. The figure compares the interaction abundance in 'evolution' and 'frozen coevolution' scenarios in relation to the 'continuous coevolution' baseline. Abundance was calculated as the number of patches harbouring each interaction as a fraction of non-destroyed patches in the landscape. Positive values indicate that the abundance in 'evolution' or 'frozen coevolution' simulations is greater than in the 'continuous coevolution' simulations, at the same fraction of habitat loss. Panels correspond to different networks in the same order as listed in Table S1. In the 'evolution' simulations, species' traits are set to their environmental optima. In the 'frozen coevolution' simulations, species' traits are set to their optimum values when the entire metanetwork is present (see Table 2).

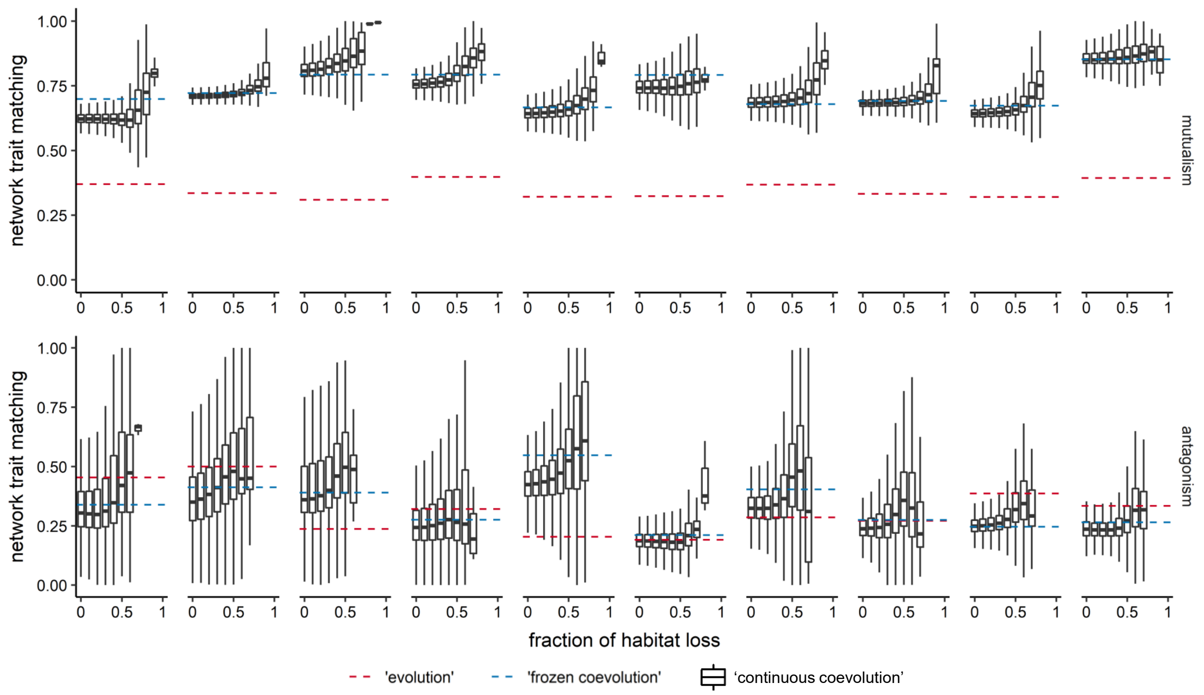


Figure S7: Network trait matching across a gradient of habitat destruction in simulations with mutualistic (top) and antagonistic (bottom) networks. Box plots summarise trait matching in all local networks in the landscape at each fraction of habitat loss in 'continuous coevolution' simulations. The red dashed horizontal lines correspond to the mean matching of environmental optima of all interacting species ('evolution' simulations). The blue dashed horizontal lines indicate the mean matching of optimum trait values when the entire empirical metanetwork is present ('frozen coevolution' simulations). Panels correspond to different networks in the same order as listed in Table S1.

**Effect of $m$**

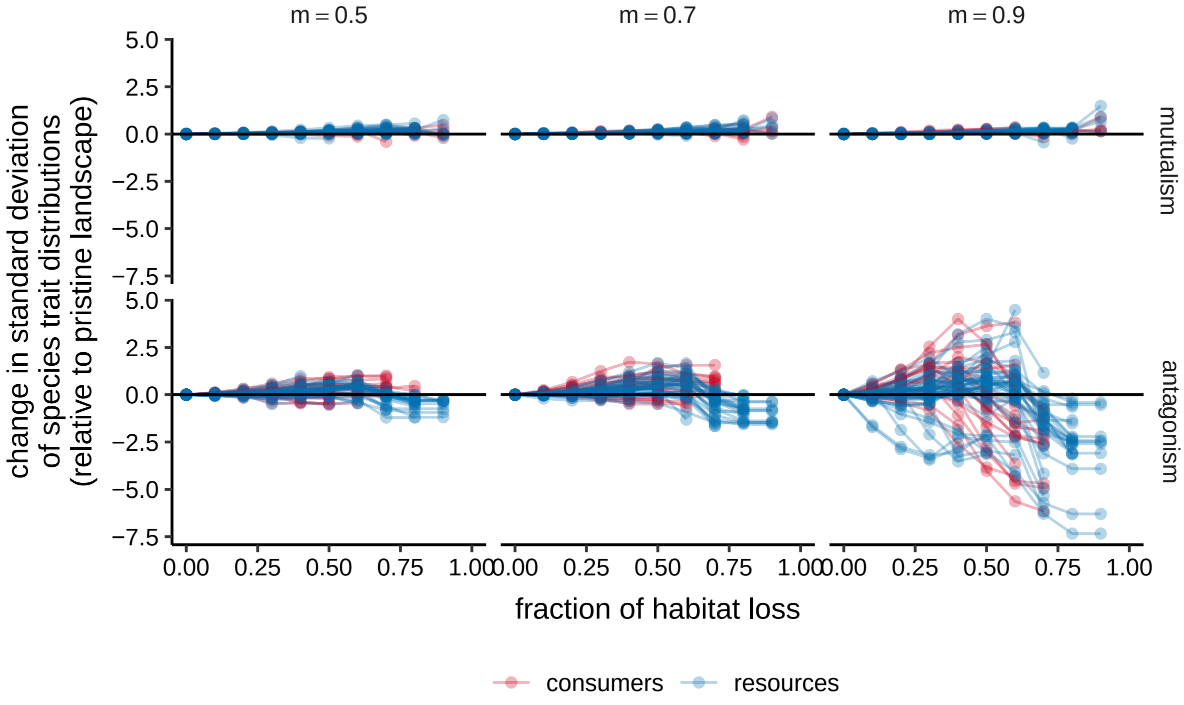


Figure S8: Change in trait variability in the landscape across an increasing fraction of habitat loss in simulations with a mutualistic (M_SD_012, top) and an antagonistic (A_PH_004, bottom) network, and different values of $m$. The change is calculated relative to the standard deviation of trait values in a pristine landscape, such that positive values indicate an increase in trait variability. Each line represents a species.

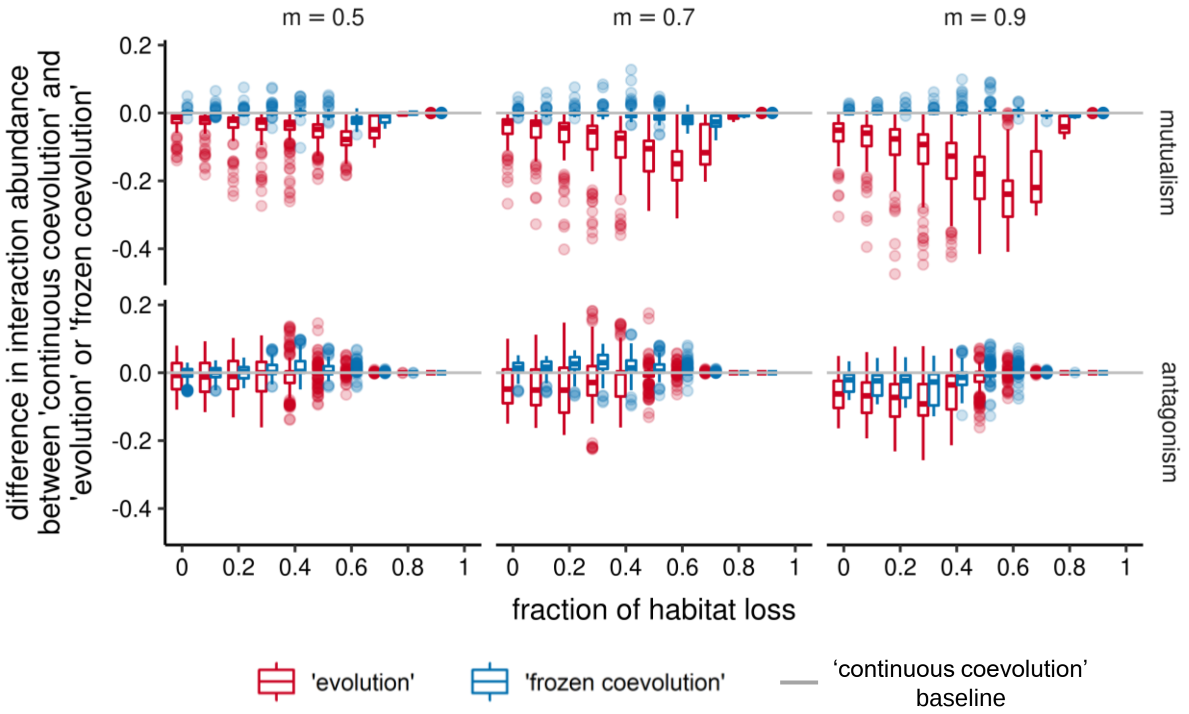


Figure S9: Relative abundance of each interaction in a mutualistic (M_SD_012, top) and an antagonistic (A_PH_004, bottom) network as habitat is progressively destroyed, in simulations with different values of $m$. The figure compares the interaction abundance in 'evolution' and 'frozen coevolution' scenarios in relation to the 'continuous coevolution' baseline. Abundance was calculated as the number of patches harbouring each interaction as a fraction of non-destroyed patches in the landscape. Positive values indicate that the abundance in 'evolution' or 'frozen coevolution' simulations is greater than in the 'continuous coevolution' simulations, at the same fraction of habitat loss. In the 'evolution' simulations, species' traits are set to their environmental optima. In the 'frozen coevolution' simulations, species' traits are set to their optimum values when the entire metanetwork is present (see Table 2).

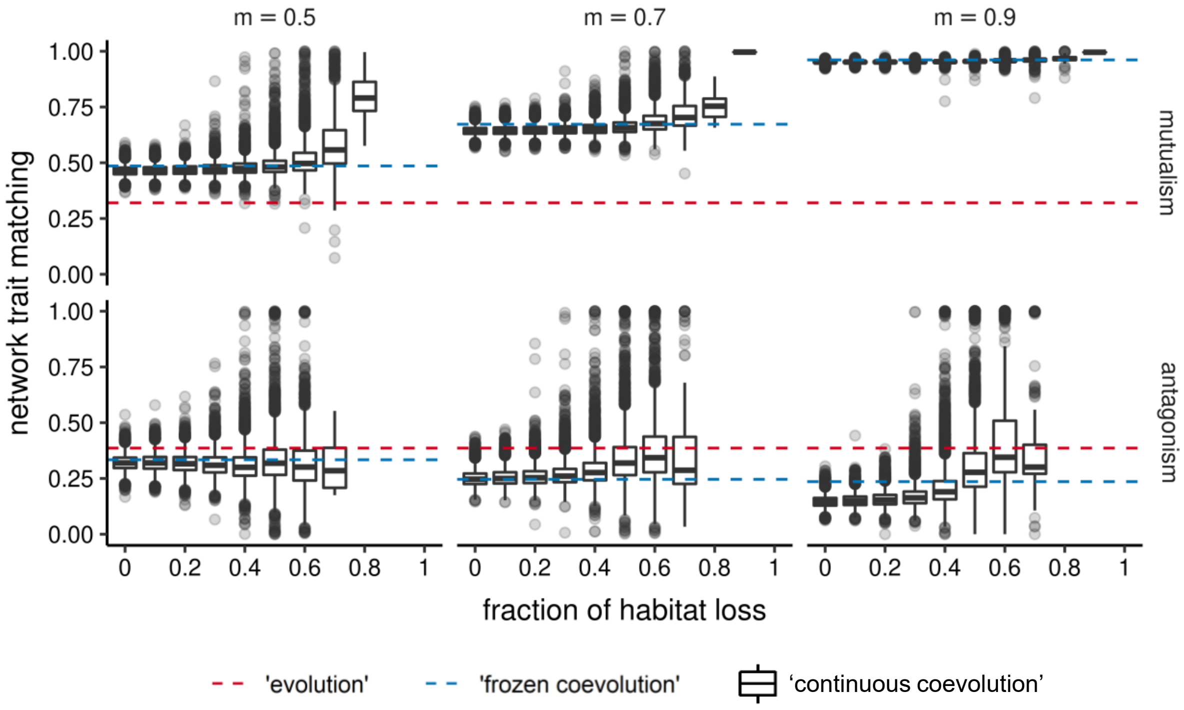


Figure S10: Network trait matching across a gradient of habitat destruction in simulations with a mutualistic (M_SD_012, top) and an antagonistic (A_PH_004, bottom) network, and different values of $m$. Box plots summarise trait matching in all local networks in the landscape at each fraction of habitat loss in ‘continuous coevolution’ simulations. The red dashed horizontal lines correspond to the mean matching of environmental optima of all interacting species (‘evolution’ simulations). The blue dashed horizontal lines indicate the mean matching of optimum trait values when the entire empirical metanetwork is present (‘frozen coevolution’ simulations).

### Effect of $\alpha$

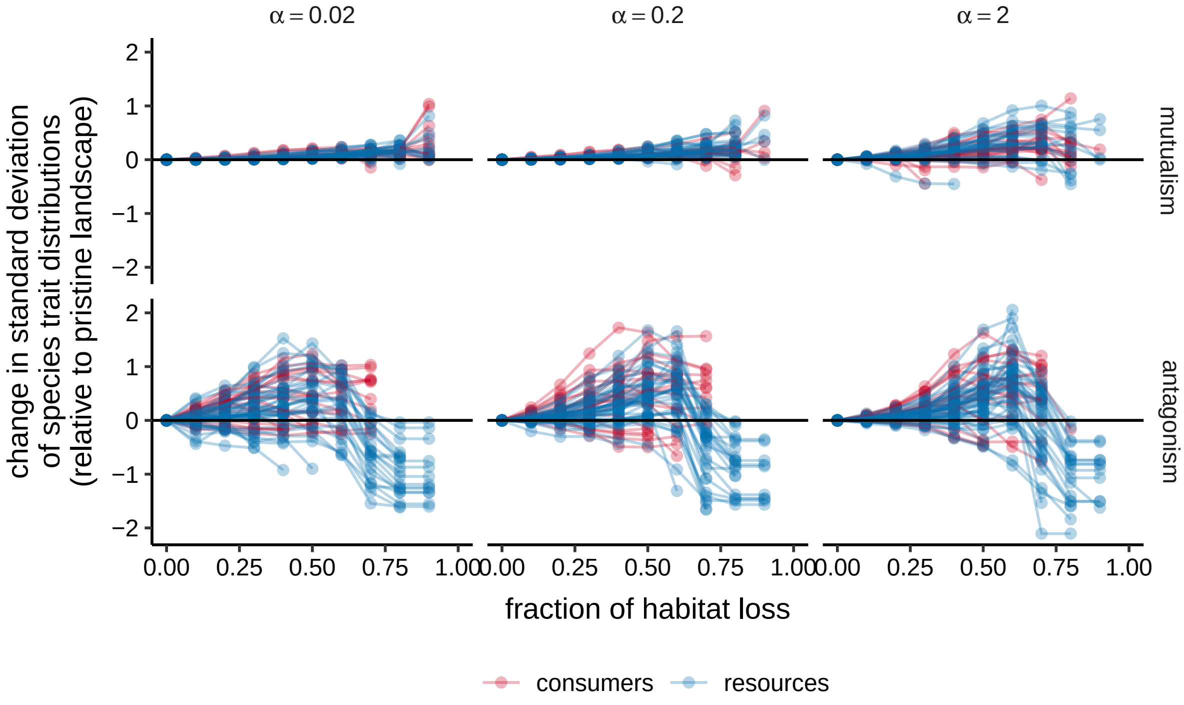


Figure S11: Change in trait variability in the landscape across an increasing fraction of habitat loss in simulations with a mutualistic (M_SD_012, top) and an antagonistic (A_PH_004, bottom) network, and different values of $\alpha$. The change is calculated relative to the standard deviation of trait values in a pristine landscape, such that positive values indicate an increase in trait variability. Each line represents a species.

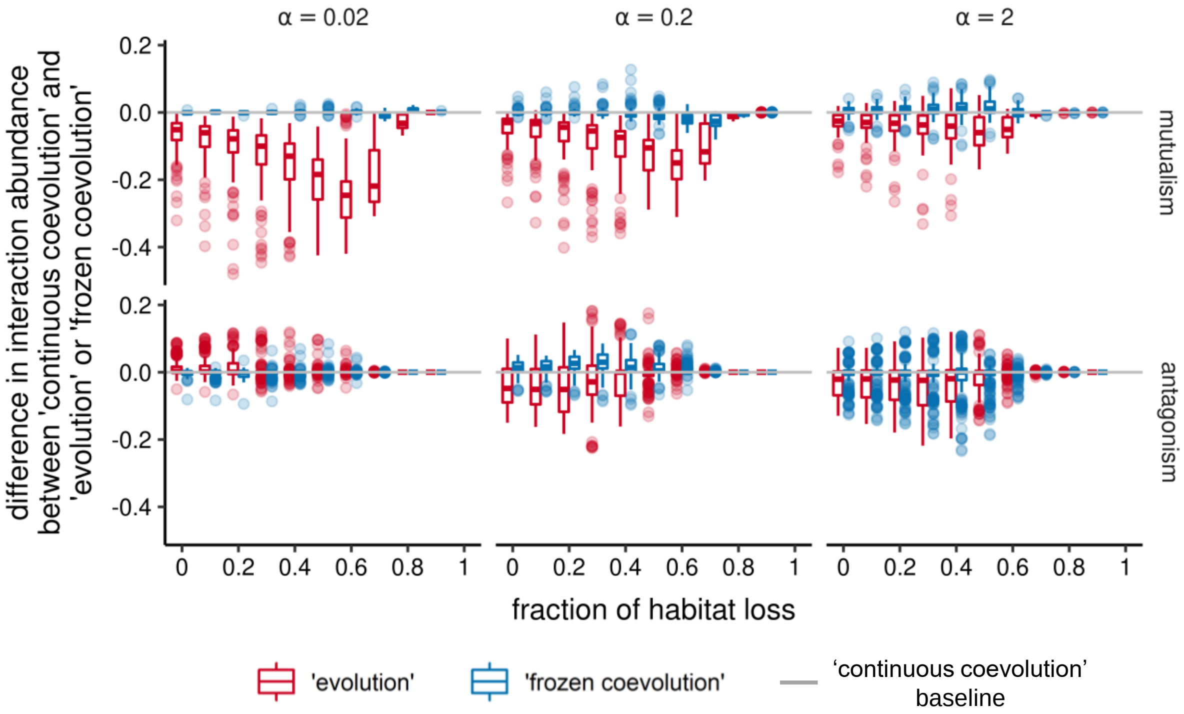


Figure S12: Relative abundance of each interaction in a mutualistic(M_SD_012, top) and an antagonistic (A_PH_004, bottom) network as habitat is progressively destroyed, in simulations with different values of $\alpha$. The figure compares the interaction abundance in 'evolution' and 'frozen coevolution' scenarios in relation to the 'continuous coevolution' baseline. Abundance was calculated as the number of patches harbouring each interaction as a fraction of non-destroyed patches in the landscape. Positive values indicate that the abundance in 'evolution' or 'frozen coevolution' simulations is greater than in the 'continuous coevolution' simulations, at the same fraction of habitat loss. In the 'evolution' simulations, species' traits are set to their environmental optima. In the 'frozen coevolution' simulations, species' traits are set to their optimum values when the entire metanetwork is present (see Table 2).

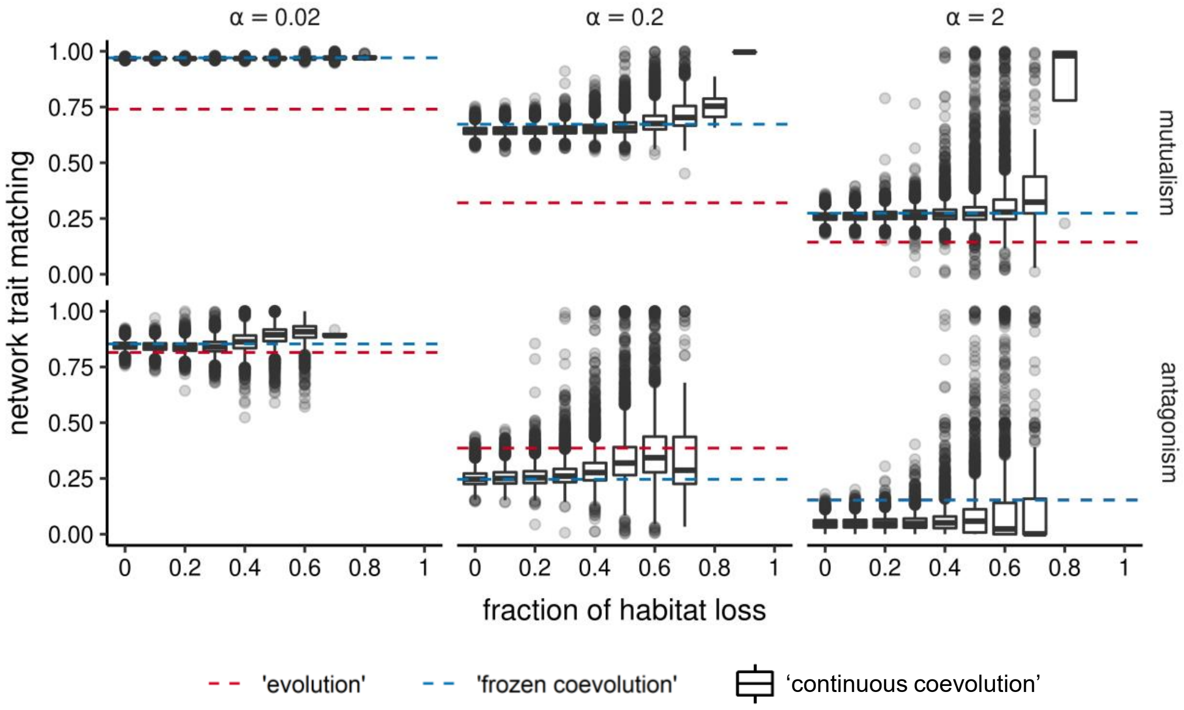


Figure S13: Network trait matching across a gradient of habitat destruction in simulations with a mutualistic (M_SD_012, top) and an antagonistic (A_PH_004, bottom) network, and different values of $\alpha$. Box plots summarise trait matching in all local networks in the landscape at each fraction of habitat loss in 'continuous coevolution' simulations. The red dashed horizontal lines correspond to the mean matching of environmental optima of all interacting species ('evolution' simulations). The blue dashed horizontal lines indicate the mean matching of optimum trait values when the entire empirical metanetwork is present ('frozen coevolution' simulations).

### Effect of $\varepsilon$

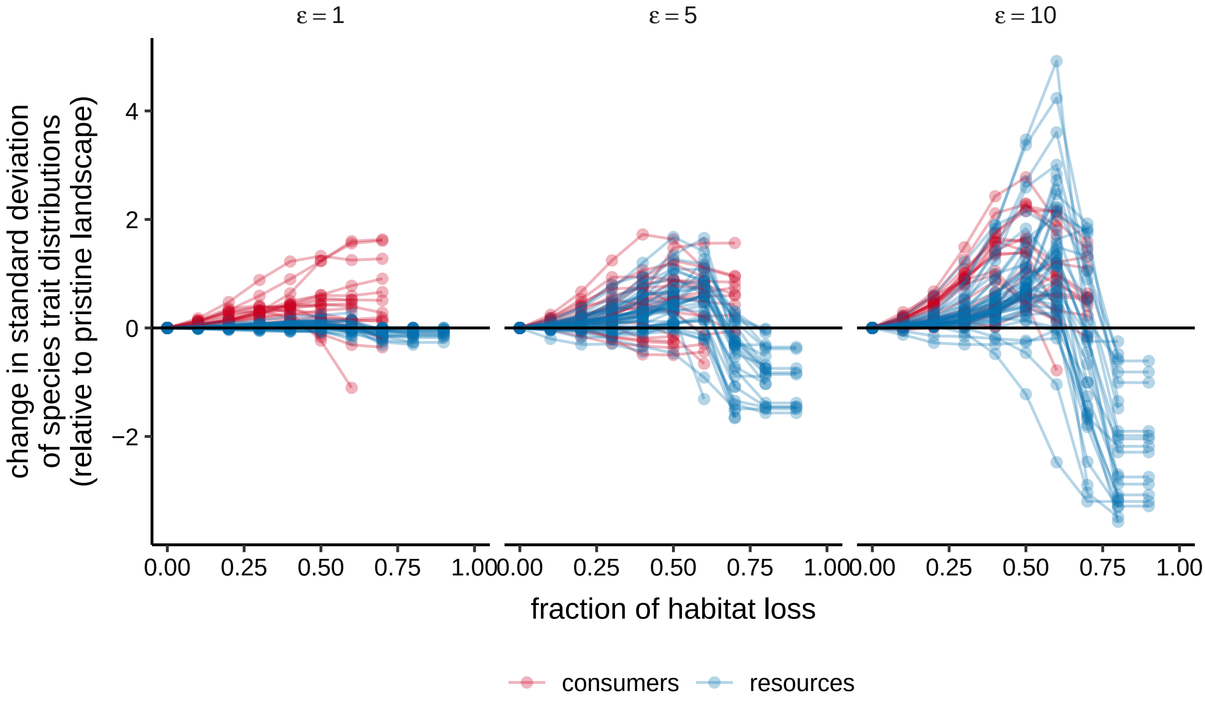


Figure S14: Change in trait variability in the landscape across an increasing fraction of habitat loss in simulations with an antagonistic (A_PH_004) network, and different values of $\varepsilon$. The change is calculated relative to the standard deviation of trait values in a pristine landscape, such that positive values indicate an increase in trait variability. Each line represents a species.

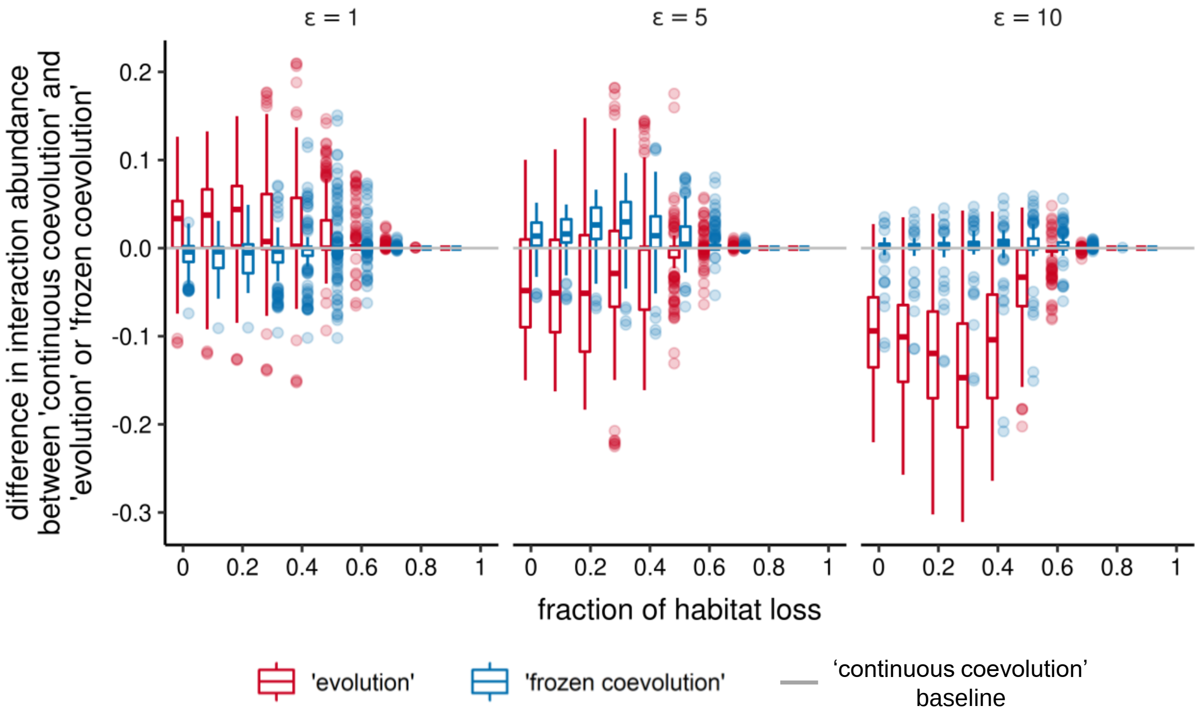


Figure S15: Relative abundance of each interaction in an antagonistic network (A_PH_004) as habitat is progressively destroyed, in simulations with different values of $\varepsilon$. The figure compares the interaction abundance in 'evolution' and 'frozen coevolution' scenarios in relation to the 'continuous coevolution' baseline. Abundance was calculated as the number of patches harbouring each interaction as a fraction of non-destroyed patches in the landscape. Positive values indicate that the abundance in 'evolution' or 'frozen coevolution' simulations is greater than in the 'continuous coevolution' simulations, at the same fraction of habitat loss. In the 'evolution' simulations, species' traits are set to their environmental optima. In the 'frozen coevolution' simulations, species' traits are set to their optimum values when the entire metanetwork is present (see Table 2).

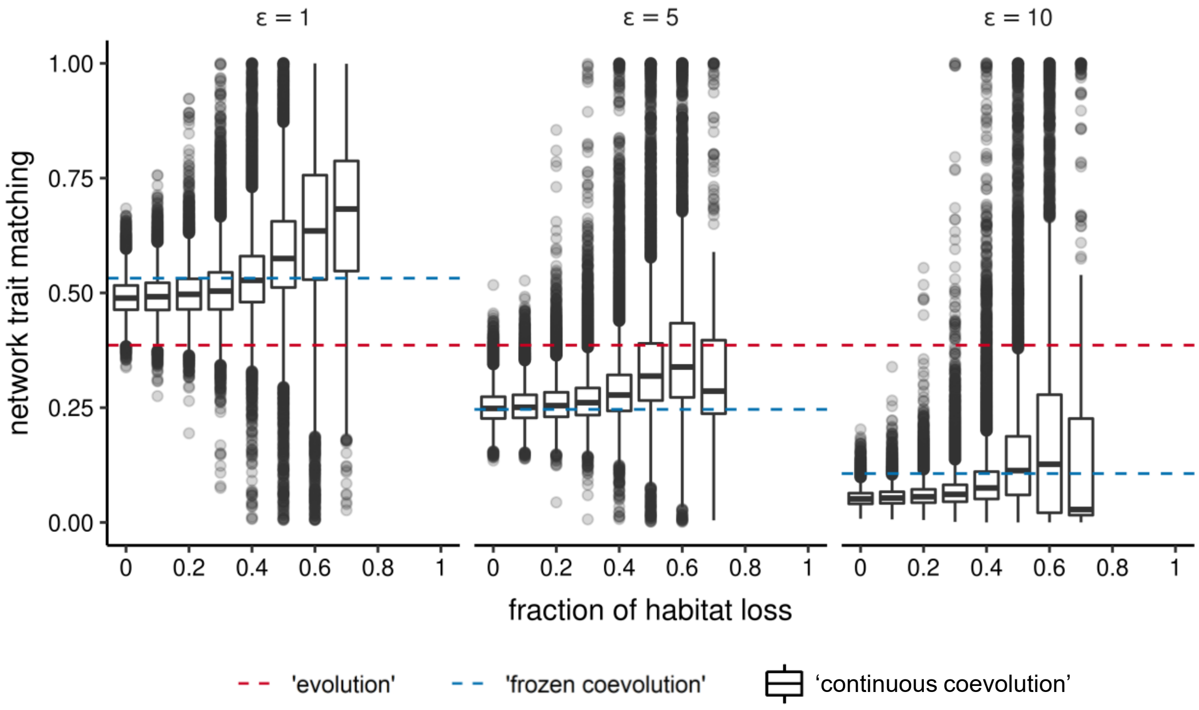


Figure S16: Network trait matching across a gradient of habitat destruction in simulations with an antagonistic (A_PH_004) network, and different values of $\varepsilon$. Box plots summarise trait matching in all local networks in the landscape at each fraction of habitat loss in 'continuous coevolution' simulations. The red dashed horizontal lines correspond to the mean matching of environmental optima of all interacting species ('evolution' simulations). The blue dashed horizontal lines indicate the mean matching of optimum trait values when the entire empirical metanetwork is present ('frozen coevolution' simulations).

### Non-random habitat destruction

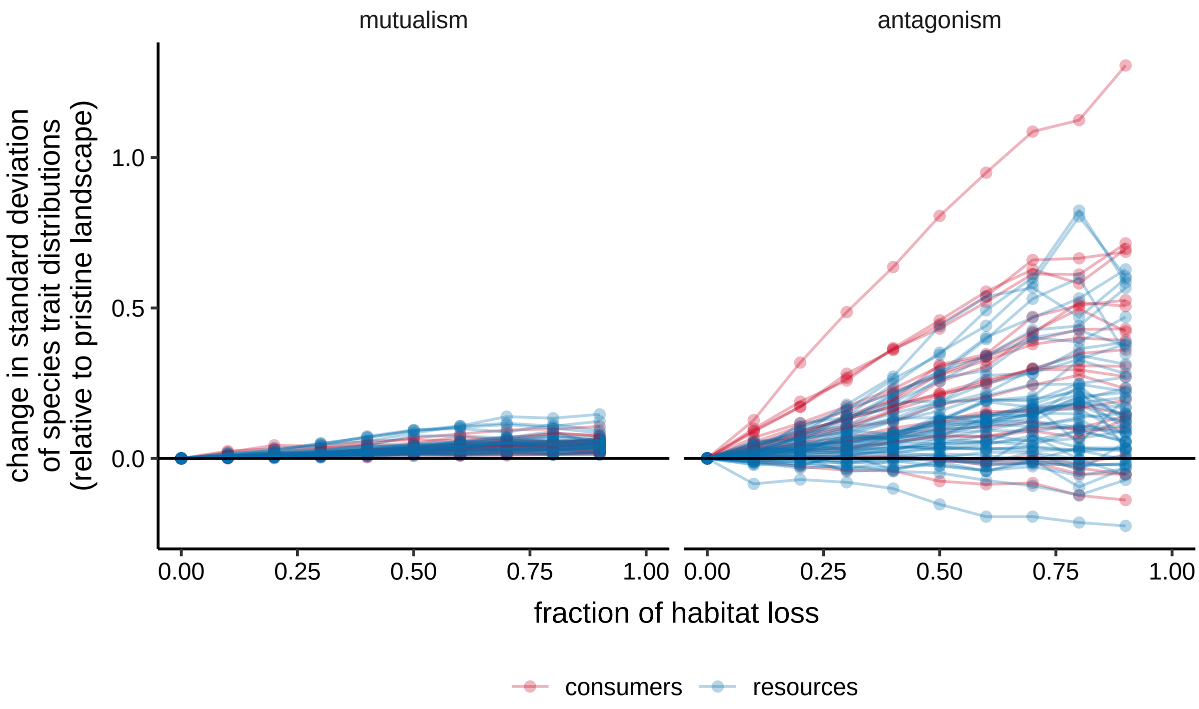


Figure S17: Change in trait variability in the landscape across an increasing fraction of habitat loss in simulations with a mutualistic (M_SD_012, left) and an antagonistic (A_PH_004, right) network, and non-random habitat destruction. The change is calculated relative to the standard deviation of trait values in a pristine landscape, such that positive values indicate an increase in trait variability. Each line represents a species.

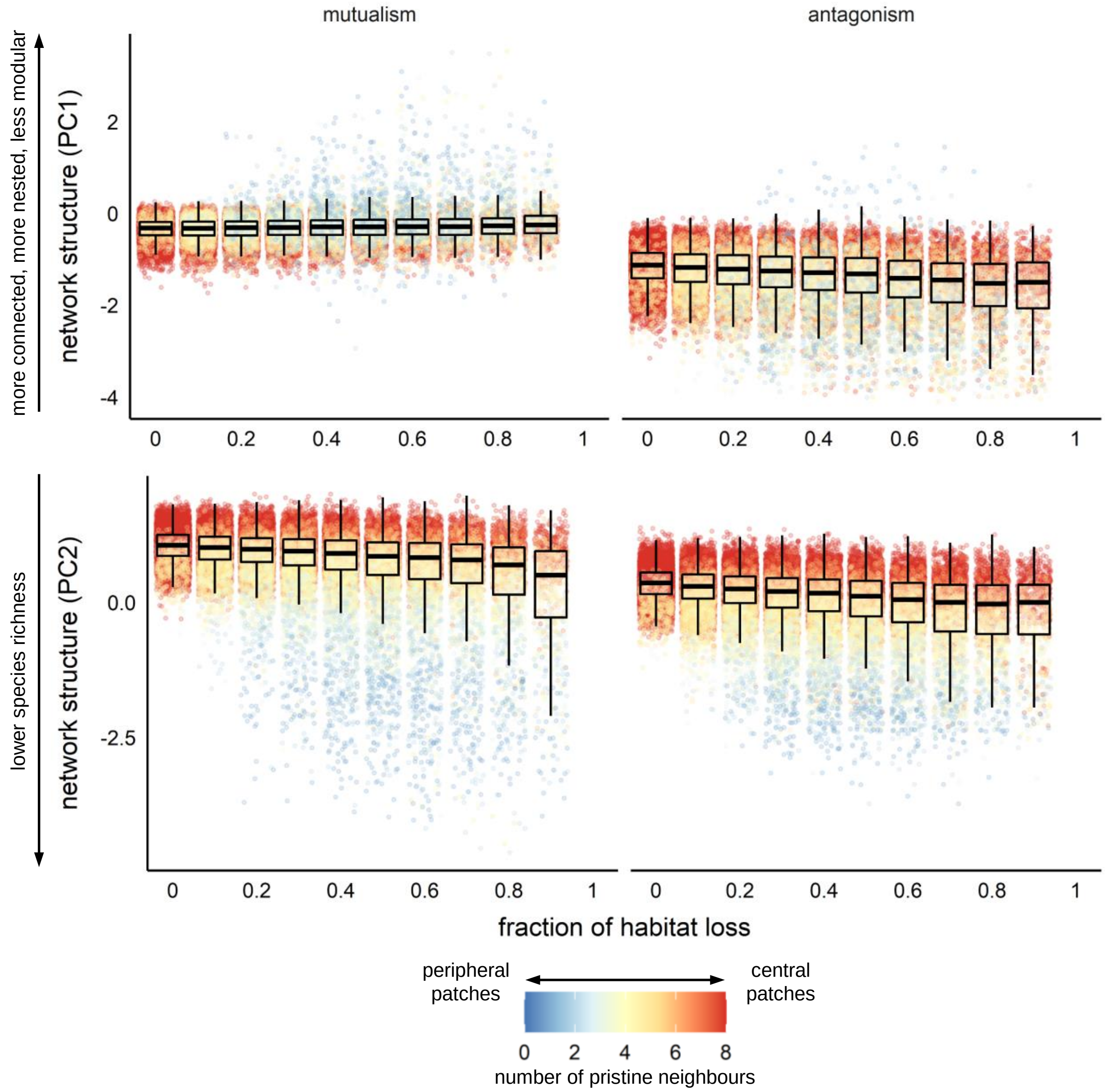


Figure S18: Effect of habitat loss on our measure of network structure of a mutualistic (M_SD_012, left) and an antagonistic (A_PH_004, right) network in simulations with non-random habitat destruction. Each point corresponds to the local network inside each patch. The colours indicate the number of adjacent pristine patches (i.e., between 0 and 8). PC1 explained 69% of variance and was strongly correlated with connectance, nestedness and modularity, whereas PC2 explained 22% of variance and was strongly correlated with network size.

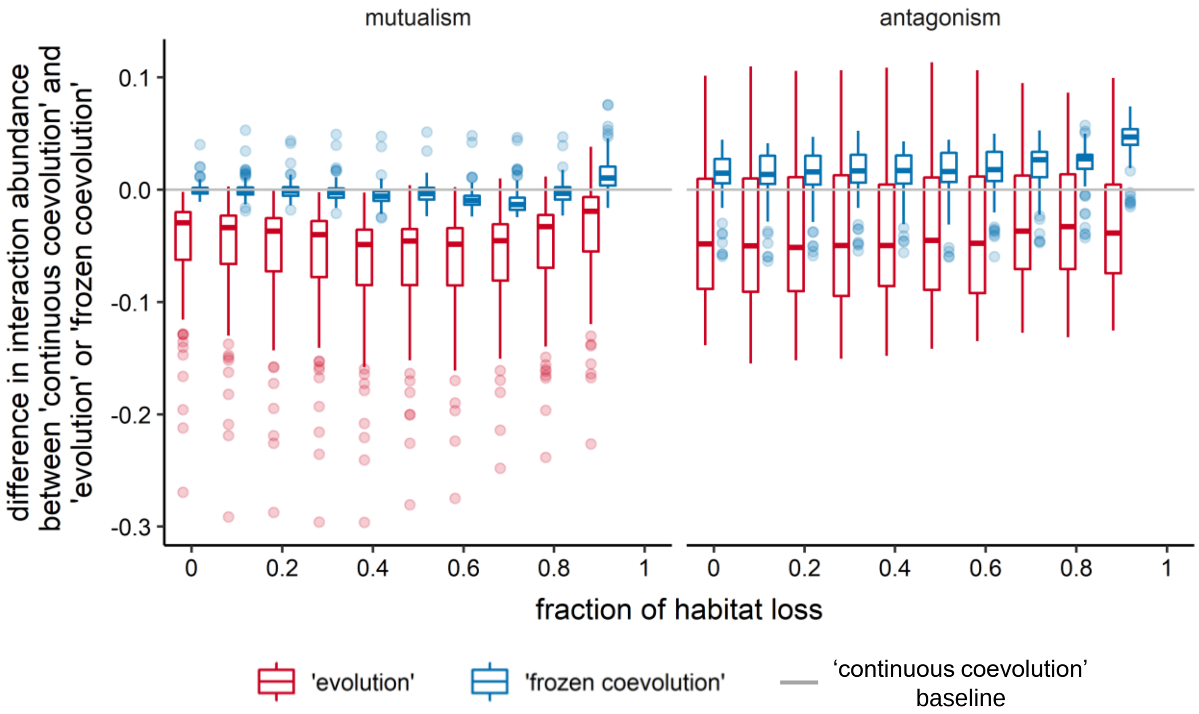


Figure S19: Relative abundance of each interaction in a mutualistic (M_SD_012, left) and an antagonistic (A_PH_004, right) network as habitat is progressively destroyed in simulations with non-random habitat destruction. The figure compares the interaction abundance in 'evolution' and 'frozen coevolution' scenarios in relation to the 'continuous coevolution' baseline. Abundance was calculated as the number of patches harbouring each interaction as a fraction of non-destroyed patches in the landscape. Positive values indicate that the abundance in 'evolution' or 'frozen coevolution' simulations is greater than in the 'continuous coevolution' simulations, at the same fraction of habitat loss. In the 'evolution' simulations, species' traits are set to their environmental optima. In the 'frozen coevolution' simulations, species' traits are set to their optimum values when the entire metanetwork is present (see Table 2).

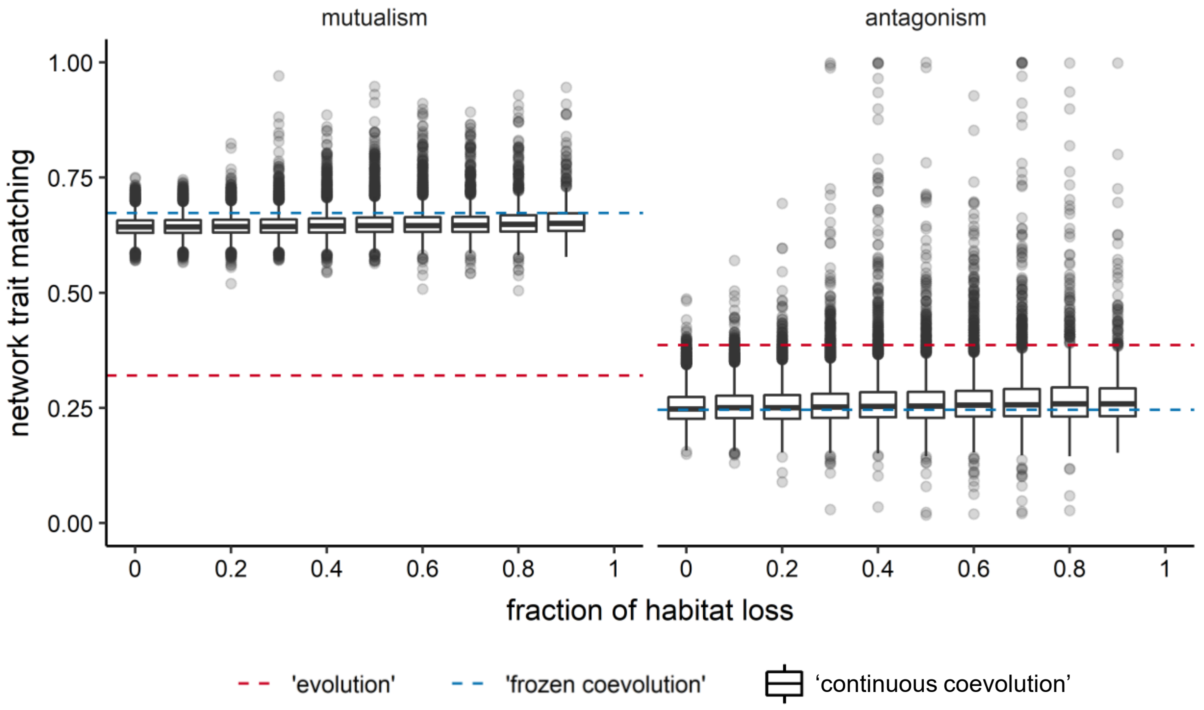


Figure S20: Network trait matching across a gradient of habitat destruction in simulations with a mutualistic (M_SD_012, left) and an antagonistic (A_PH_004, right) network, and non-random habitat destruction. Box plots summarise trait matching in all local networks in the landscape at each fraction of habitat loss in 'continuous coevolution' simulations. The red dashed horizontal lines correspond to the mean matching of environmental optima of all interacting species ('evolution' simulations). The blue dashed horizontal lines indicate the mean matching of optimum trait values when the entire empirical metanetwork is present ('frozen coevolution' simulations).
